# Helical motors and formins synergize to compact chiral filopodial bundles: a theoretical perspective

**DOI:** 10.1101/2023.07.24.550422

**Authors:** Ondrej Maxian, Alex Mogilner

## Abstract

Chiral actin bundles have been shown to play an important role in cell dynamics, but our understanding of the molecular mechanisms which combine to generate chirality remains incomplete. We numerically simulate a crosslinked filopodial bundle under the actions of helical myosin motors and/or formins and examine the collective buckling and twisting of the actin bundle. We find that the myosin spinning action effectively “braids” the bundle, compacting it, generating buckling, and enhancing crosslinking. Stochastic fluctuations of actin polymerization rates also contribute to filament buckling and bending of the bundle. Faster turnover of transient crosslinks attenuates the buckling and enhances coiling and compaction of the bundle. Formin twisting action by itself is not effective in inducing filopodial coiling and compaction, but co-rotating formins synergize with helical motors to coil and compact the actin bundle. We discuss implications of our findings for mechanisms of cytoskeletal chirality.

## 1. Introduction

The cytoskeleton shapes cells, tissues and organs (Fletcher and Mullins, 2010). Actin filaments are crucial elements of the cytoskeleton, and are polar and chiral right-handed helices with distinct plus and minus ends (Jegou and Romet-Lemonne, 2020). From physiology, it is clear that the molecular polarity, helicity and chirality of actin filaments propagate to cytoskeletal arrays, cells, and tissues, which ultimately underlie important left-right asymmetries in development and complex organisms (Inaki et al., 2016). However, the mechanisms of this propagation remain largely unclear.

Two mechanical molecular processes were proposed to be the starting points for chirality generation in the cytoskeleton (Naganathan et al., 2016; Jegou and Romet-Lemonne, 2020). The first one is based on the action of formin, the “leaky capper” protein that stays on the growing plus end of the filament, thereby mediating the process of actin polymerization (Fig. 1A). Because of actin’s helical structure, formin can rotate around the filament axis in the process of growth, or, if formin is fixed to a larger structure, an actin filament could rotate by being “extruded” from formin (Mizuno et al., 2011), in which case formin effectively applies a torque to the filament. The second symmetry-breaking process could be triggered by myosin molecular motors that in some cases move helically around actin filaments (Nishizaka et al., 1993), effectively applying both torque and force. Our goal here is to computationally examine how these two plausible pathways of chirality emergence act in a small polar actin bundle.

**Figure 1:**
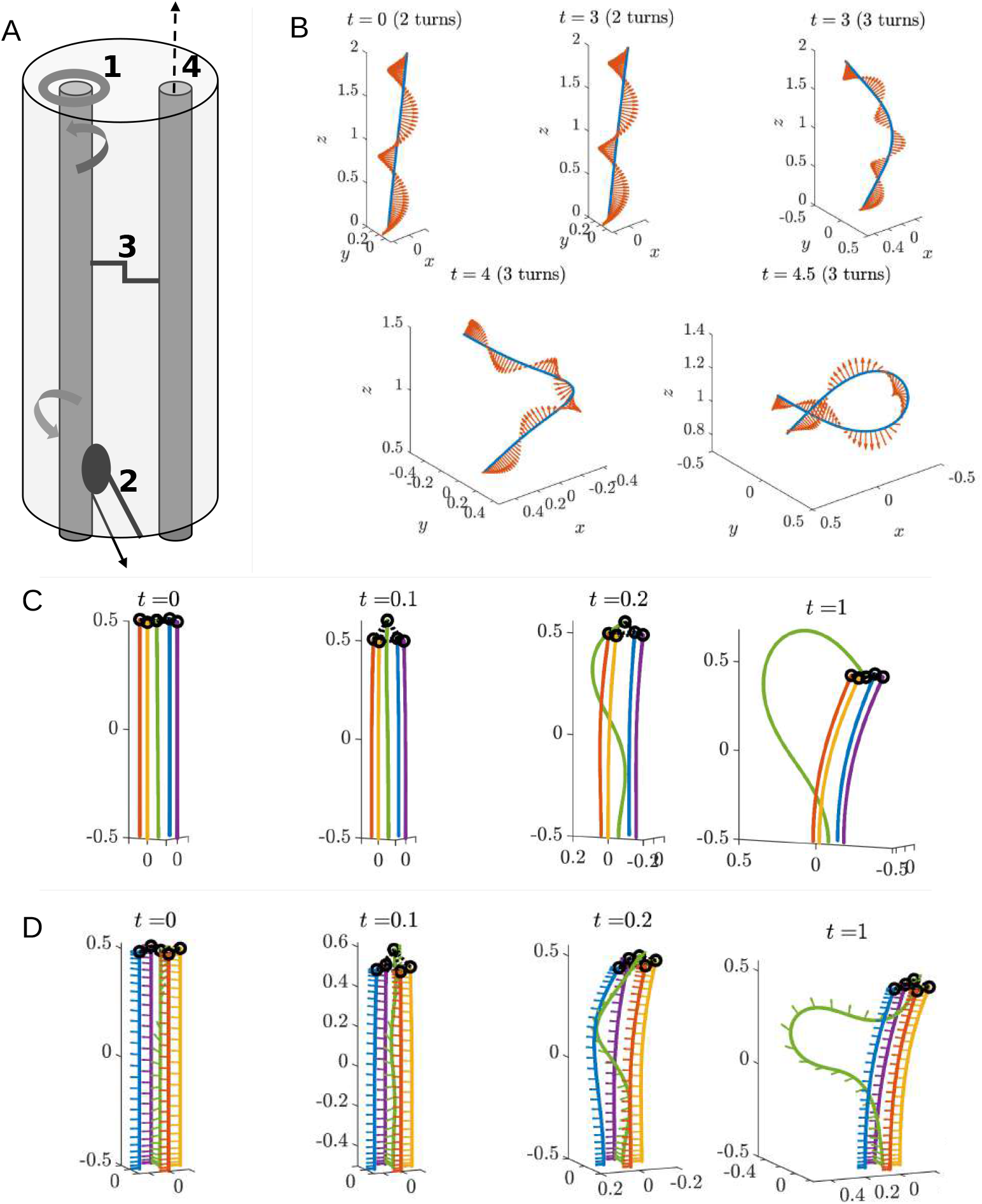
Actin supercoiling and its effect on polymerization-induced buckling. (A) Mechanics of actin filaments (two thin dark rods) inside the filopodial shaft (transparent cylinder). (1) Formins (doughnut-shaped) generate effective torque applied to filament plus ends (at the top). (2) Myosin V motors, attached to the filopodial shaft, move helically toward the filament plus ends, thereby applying a force which both pulls the filaments toward their base and spins the filaments around the filopodial shaft. (3) Transient crosslinks interconnect the filaments. (4) Filaments polymerize from their endpoint with dynamic random rates. (B) Simulation of a supercoiling actin filament. In this simulation, formin acting at one end of the filament applies a torque and fixes an effective twist which is constant in time. The blue line shows the fiber position, and the red arrows show material frame vectors (which demonstrate the twist). We consider a simulation where the material frame is twisted in two rotations from start to endpoint, where perturbations relax to straight, and another where the material frame is twisted in three rotations, where perturbations grow in time and supercoiling is observed. (C) Buckling in a five-filament bundle due to the polymerization of the central (green) filament. The polymerization is 1 *μ*m/s on 0 ≤ *t* ≤ 1 s. Filaments are color-coded for visualization. The tips of the outer filaments are permanently crosslinked (dashed segments) to the tip of the central filament; rings are the endpoints of the crosslinks. For time points after *t* = 1, see Fig. S1. (D) Buckling in a five-filament bundle due to the polymerization of the central (green) filament, with the increased twist (*N*_*L*_ = 0.5 pN ·*μ*m) added along the central filament. The polymerization is 1 *μ*m/s on 0 ≤ *t* ≤ 1 s. Arrows show the material frame vectors.

There is no consensus yet on the chiral action of formins. On the one hand, Mizuno et al. observed actin filaments rotating clockwise (CW), if viewed from the plus end, when growing with a formin on the plus end (Mizuno et al., 2011). This would imply a formin-generated CW torque on the filament plus end. On the other hand, no supercoiling was observed when long actin filaments grew from formins attached to a substrate (Kovar and Pollard, 2004). If there is a formin-generated torque, then a sufficiently long filament must supercoil, so this result implies no or insignificantly weak torque. To explain this paradox, Shemesh et al. (Shemesh et al., 2005) proposed a mechanism where formins twist actin filaments until this twist is no longer energetically favorable, at which time a “screw” rotation dissipates the torsional strain. This model was expanded in the recent study of Li and Chen (Li and Chen, 2022), which considered it in tandem with two other possible models of formin-actin interaction. The conclusion of that study is that, for some mechanisms, the formin can apply a rapidly fluctuating torque on the filament end, and that the net torque would be on the order of 100 pN·nm (Li and Chen, 2022). Moreover, the Li and Chen model (Li and Chen, 2022) predicts that the torque can be both CW and counterclockwise (CCW).

Several reports pointed out that some myosin motors are helical, walking around actin filaments along a helical path, so when the motors are attached to a larger structure, they not only exert a force on a filament, but also apply torque to it. Tanaka et al. reported that muscle myosin could be either a CW or a CCW motor (Tanaka et al., 1992), while a later study proposed it to be exclusively CW motor (Nishizaka et al., 1993). (In the literature, the terms “right-” and “left-handed”’ are often used for the helical motors; here, to avoid confusion, we call them CW and CCW (rotating the filament CW and CCW if viewed from plus end), respectively. Our view direction will always be from the plus end when mentioning CW or CCW rotation, and so we will skip mentioning the point of view henceforth.) Myosin V was reported to be a CCW spiral motor (Cheney et al., 1993; Ali et al., 2002). Myosin X was shown to be the CCW helical motor (Sun et al., 2010). Myosin 1D was found to be a CCW motor (Lebreton et al., 2018).

These formin and myosin actions are relevant for actin bundle dynamics in filopodia, which are finger-like bundles which protrude from the cell edge and measure several to tens of microns in length and tenths of microns in diameter (Jacquemet et al., 2015). They consist of a parallel bundle of 10 to 30 actin filaments (Jacquemet et al., 2015) crosslinked by fascin (Aratyn et al., 2007) and other crosslinkers. The filaments polymerize and grow at the plus ends at the filopodial tip (Mallavarapu and Mitchison, 1999), while the bundle undergoes retrograde flow (away from the tip) driven by myosins at the filopodial base (Forscher and Smith, 1988; Aratyn et al., 2007). Filopodia play a variety of mechanical and sensory functions (Davenport et al., 1993) and are implicated in wound healing (Wood et al., 2002), cancerogenesis (Arjonen et al., 2011) and development (Fierro-González et al., 2013).

Several examples of chiral filopodial behavior have been reported. For example, it was shown that actin filaments in filopodia flow toward the cell body, rotating in the CCW direction (Forscher and Smith, 1988). Individual filopodia were observed to rotate around their longitudinal axes in the CCW direction (Tamada et al., 2010), in a myosin V dependent way. Adherent filopodia, formed during the spreading of cells, tended to change the direction of their extension in a chiral fashion, acquiring a left-bent shape, if viewed from above, in a myosin X- and formin-dependent way (Li et al., 2023). Most notably, not only chiral movements, but also buckling and CCW helical rotation, as well as twisting and coiling of the filopodial actin bundle were reported (Leijnse et al., 2022). These deformations were myosin-, but not formin-dependent (Leijnse et al., 2022).

In filopodia, formins are at the tip, capping and elongating filaments’ plus ends (Mellor, 2010). Formins there are a part of the “tip complex,” a dense large protein patch that may or may not be relatively rigid (Mellor, 2010), so there is a possibility that formins are mechanically restrained and apply a net torque to the growing filament ends. In our model, we will examine the possible CCW formin-generated net torque (Fig. 1A), and then also consider changes in model predictions when the torque is CW. Various types of myosin motors are found at different parts of filopodia (Leijnse et al., 2022). For example, there is abundant expression of myosin V at the base of filopodia (Leijnse et al., 2022). It is reasonable to assume that helical myosin motors there are fixed to relatively rigid structures near the filopodial base, and if these motors are not very processive, their powerstrokes that are oriented diagonally relative to the filament centerlines have components along the circumference of the filopodial bundle that rotate, or spin, the filaments around the filopodial axis. In the model, we will consider only CCW myosin-generated net torque (Fig. 1A), since we need only consider the direction of the myosin torque *relative* to that of formin (for which we consider both directions).

The general questions we wish to answer are: how do the chiral molecular mechanics of individual filament/myosin/formin actors play out on the scale of cytoskeletal arrays, creating collective chirality of actin filaments? Would not crosslinking interfere with collective chirality (one would imagine that stapling two filaments together prevents their rotation)? And, do myosins and formins cooperate or inhibit each other’s actions? To address these questions, we numerically simulate a crosslinked filopodial bundle under chiral myosin and/or formin actions and quantitatively examine the collective buckling and coiling response. We find that the myosin spinning action effectively “braids” the actin bundle, compacting it, generating buckling, and enhancing the crosslinking. Stochastic fluctuations of actin polymerization rates also contribute favorably to filament buckling and bending of the bundle. Faster turnover of transient crosslinks removes the constraints that give buckling, thereby attenuating it, but at the same time enhances coiling and compaction of the bundle. Formin-generated twisting is much less effective than myosin-generated spinning in inducing filopodial coiling; however, with myosin activity, co-rotating formins are essential to give proper bundle compaction.

## 2. Materials and Methods: computational model

In a recent combined experimental-theoretical study, the emergence of chirality and twist in filopodia was modeled by considering the actin bundle as a continuum active polar gel (Leijnse et al., 2022). In our case, the relatively small number of filaments and associated molecules in filopodia makes a discrete approach, in which filaments are treated as elastic rods under the action of motor forces, more appropriate. Successful mathematical analyses of deformations of bent and twisted rods (Stump, 2000) and buckling of actin filaments (Berro et al., 2007) and bundles (Martiel et al., 2020) allow us to build a detailed mechanical model of a dynamic filopodial bundle. Here, we explain the computational model we used to address the questions raised in the Introduction. For the sake of biologist readers, we explain the model mostly in words; the details are described in the Supplemental Material (SM). A schematic of the model is also shown in Fig. 1A.

### Actin filament mechanics

We adopt our published model (Maxian et al., 2022) to describe actin filaments as cylindrical, slender, inextensible elastic rods undergoing and resisting bending and twisting deformations. The filaments, which we sometimes call fibers, move, rotate and deform under the influence of the following forces and torques:

- Elastic bending force tends to straighten the filament’s centerline.
- Elastic twisting force tends to minimize the filament’s twist (relative rotation of neighboring cross sections of the rod around the centerline).
- Viscous resistance from the movement of the filament through the fluid it is immersed in.
- Elastic force between neighboring filaments generated by deformations of the crosslinks that couple pairs of filaments.
- Steric force between pairs of filaments that are in touch with each other, preventing one filament from passing through another.
- Active force generated by myosin motors, which we will call motor force for brevity.
- Active torques generated by formins.

Mathematics of the elastic bending and twisting forces and of the viscous resistance force are explained in the SM. Below, we qualitatively explain the boundary conditions for the filaments, model geometry, active motor and formin forces and torques, and crosslinking and steric forces. Then, we discuss actin polymerization, and how we quantify the filopodial bundle’s shape.

### Boundary conditions

The minus ends of the filaments always remain at the base, *z* = 0, so we neglect possible retrograde flow of the filaments and consider them cantilevered into a solid surface of the actin cortex of the cell. Each minus end is clamped at the base, so both its position and orientation, as well as the orientation of the filament cross section there, do not change in time. The plus ends are free to move and twist (in the precise mathematical sense explained in SM).

### Model geometry

To build intuition about filopodial mechanics, we first consider a small five-filament bundle in which there is a central filament and four parallel peripheral filaments, the minus ends of which are located on the vertices of a square (Fig. 1C,D, 2B, 3A,B). The filaments are spaced so that distances between pairs is approximately equal to the crosslink rest length. Then, we consider a larger 22-fiber bundle (realistic number for a filopodium) arranged into four rings (1, 3, 6, and 12 filaments) (Fig. 4). The rings are spaced 100 nm (the crosslink rest length) apart, and fibers are positioned with uniform spacing around each ring. Thus for the outermost ring, the 12 fibers are positioned at ***X***(*s*) = (3*ℓ*_*c*_ cos (2*πi/n*), 3*ℓ*_*c*_ sin (2*πi/n*), *s*) for *i* = 0, … 11, where *ℓ*_*c*_ is the crosslink rest length (100 nm).

### Motor forces

As shown in Fig. 1A and discussed in the Introduction, a helical myosin motor generates power strokes along a helical path on the cylindrical surface of a filament. The resulting force has two components, one of which tries to move the filament retrogradely toward the filopodial base, which we assume is balanced by the reaction of the cell cortex at the base (Evans et al., 1997). Another component is directed along the circumference of the filopodium, so that this force spins the filaments around the long axis of the filopodium. Thus, the motors push the filaments into a circular trajectory, where the circle has radius *R* equal to the filament distance from the central filopodial axis. At a point ***x***, the force ***f*** ^(mot)^ can be computed by first converting ***x*** into cylindrical coordinates (*R, θ, z*), and using these to define a vector tangent to the circle at ***x***, via ***t*** = (− sin *θ*, cos *θ*, 0). The force density is then given by 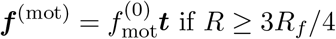 and *z* ≤ *c*_*m*_*L* (*L* is the filament length), so that the motors act only on the outer 1*/*4 and bottom *c*_*m*_ of the filopodium. (Effectively, the motors from the boundary can only reach filaments that are relatively close to the boundary. For the 5-filament bundle, these are the four outer filaments, and for the 22-filament bundle, these are the twelve filaments of the outer ring.) Two parameters here defining the motor forces include 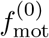, which is the magnitude of the force density in units of pN/*μ*m, and *c*_*m*_, the fraction of the filaments on which the motors act. We will vary parameter *c*_*m*_ between 1/2 (lower half of the filipodial boundary is covered with motors) and 1 (the whole length of the filipodial boundary is covered with motors). As mentioned in the Introduction, we consider motors which spin filaments in a CCW direction around the filopodium. This model implies that the cylindrical surface to which the motors are bound is rigid, which is likely an oversimplification.

### Formin and motor torque

Formins apply a constant torque only to the polymerizing plus ends, as discussed in the Introduction. There is no torque on the static ends. We first consider a torque which tries to twist the filament in a CCW direction around the centerline, in the same direction as the motor spin. We also discuss the effect of the opposite torque direction.

### Crosslinking

In the model, the crosslinks connecting filament pairs are linear springs with finite rest length whose ends connect to the filaments by completely flexible joints, so possible bending forces are ignored. Also, the crosslinks bind, effectively, to the 1D centerlines of filaments, so potential effects from the specific distribution of binding sites along the helical grooves of the filaments are also ignored. The crosslinking kinetics consist of unbinding of each crosslink with a constant, force-independent, rate, and of crosslink binding to a pair of filaments with length-dependent rate. When a crosslink unbinds, both of its ends dissociate from the filaments at once. The binding rate is a function of the distance between the two points on the filament pair to which the binding occurs, and is maximal if this distance is equal to the crosslink rest length. The detailed kinetics and thermodynamics of these processes are discussed in the SM. In addition to transient crosslinking, we also sometimes consider a filopodial protein tip complex to which the plus ends of the filaments bind. This cap is modeled by a network of permanent crosslinks interconnecting the plus ends of the filaments, whether static or polymerizing. These permanent crosslinks have the same mechanical properties as the transient ones, just different kinetics.

### Steric forces

When any two points on distinct filaments become too close, we apply a strong, short-range repulsive force that prevents them from passing through each other, while not perturbing any dynamics of the non-overlapping filaments. The details of this force can be found in the SM, and in (Maxian, 2023, Sec. 9.2.1)

### Modeling actin polymerization

Actin polymerization is modeled with elongation of a filament at its plus end with a certain rate 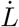. It was argued that in filopodia the polymerization is stochastic due to the relatively small size of the system and consequent large fluctuations of the actin monomer concentration (Lan and Papoian, 2008). One additional source of stochasticity could be persistently different chemical conditions of different filament plus ends, i.e. one filament’s end associated with formin, another one with VASP protein, etc. To account for that, we first (in the five-filament bundle) simulated growth rates constant in time but varying between the filaments. Then, (in the 22-filament bundles) we make the growth rate of each filament a random variable which obeys the Ornstein-Uhlenbeck process:

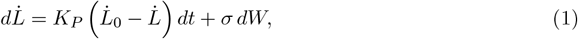

where the constant *K*_*P*_ = 2/s is chosen so that deterministic fluctuations relax to the base value 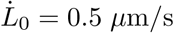 on timescales 0.5 s. The noise strength 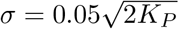 is chosen so the (stationary) standard deviation of the growth rate is 0.05 *μ*m/s. In simple words, the average rate is the same for all filaments, but each filament’s rate independently fluctuates in time with certain variance. In the model, we neglect the experimentally-observed dependence of the polymerization rate on force and torque (Yu et al., 2017). Finally, filament polymerization is capped when the length exceeds double the initial length (this also relaxes the twist deformations).

### Model parameters

The elastic and mechanical parameters characterizing actin filaments, crosslinkers, motor and formin forces and torques, polymerization rates, crosslinker kinetics and viscous drag coefficients have all been measured or estimated in experimental literature. The full list of parameters, their values and respective citations are gathered in the Supplemental Table. We vary some of the parameters, like formin torque, crosslinker kinetics and variance of the polymerization rates, to evaluate their effect on the bundle dynamics. The model predictions are relatively robust to few-fold changes in other parameters.

### Quantification of the filopodial bundle’s shape

In order to compare predicted filopodial dynamics and mechanics at different model regimes and parameters, we need to quantify the buckling and twisting of the filaments. Intuitively, we are interested in three parameters: the vertical extension of the filaments (in the direction of the bundle axis, or *z* direction), the filaments’ bend, and their twist around the bundle axis. To quantify these, we transform the coordinates of each fiber from Cartesian to cylindrical, where the origin for the polar grid is set by the closest point on the central fiber. This gives a set of coordinates (*r*(*s*), *θ*(*s*), *z*(*s*)) for each fiber. The vertical extension can then be tracked by looking at *z*(*L, t*). We also sometimes plot “the angle from straight” equal to cos^−1^((*z*(*L, t*) − *z*(*L*, 0)) */L*). The amount of twist can be quantified by the number of rotations around the axis of either the filament endpoints, (*θ*(*L, t*) −*θ*(*L*, 0))*/*2*π*, or their midpoints, (*θ*(*L/*2, *t*) − *θ*(*L/*2, 0))*/*2*π*. To study the amount of bend, we compute the curvature of the fibers (magnitude of the derivative of the tangent vector with respect to arclength) and average the curvature over the whole fiber (in the so-called *L*^2^ norm). We then normalize the curvature by the curvature of a circle of radius equal to the initial fiber length *L*(0), which is 2*π/L*(0). Together, these statistics convey how bent and twisted the filaments are. They are similar to the quantifiers used in (Grason and Bruinsma, 2007; Tee et al., 2023) to measure chirality and twist. It is also useful to plot the traces or trajectories of the fiber endpoints over time projected onto the (*x, y*) plane.

### Numerical procedure for simulating the actin bundle

At each computational step, current geometries of all fibers and engaged crosslinks are used to compute the elastic bending and twisting forces, steric and cross-linking forces, and motor and formin torques. By balancing the net forces and torques with viscous resistive forces, the linear and angular velocities of the filaments are calculated, and their geometries are updated. In addition, the increase of the filaments’ lengths by polymerization is calculated, and some of the existing crosslinks are dissociated, while new crosslinks are engaged with filament pairs according to their dynamics. Marching in small time increments from step to step evolves the bundle. Here we use a time in seconds, which is a natural time scale for the cell-scale forces and viscosities. As discussed in the SM, twist equilibrates 𝒪(1*/ϵ*^2^) faster than bending (Powers, 2010), where *ϵ* is the fiber aspect ratio. As such, in our simulations twist deformations, while technically dynamic, are typically in a steady state controlled by the amount of the formin twist (see the SM).

## 3. Results

### 3.1 High torque supercoils filaments and attenuates polymerization-induced buckling

#### 3.1.1 Actin supercoiling above a critical twist density

We first investigate the possibility that actin supercoiling (Bibeau et al., 2023) could play a role in the bending of filopodia. We start by estimating the magnitude of the twist at the plus end that supercoils an actin filament. Linear stability analysis of the model equations (see the SM) states that the critical twist, above which the filament supercoils, corresponds to 2.4 360-degree turns of the plus end. We test this prediction numerically by applying 2 turns to the plus end and observing that the filament relaxes to a straight shape, and then applying 3 turns to the plus end and observing that the filament supercoils (Fig. 1B).

According to our estimates in the SM, the characteristic formin-generated torque will result in ≈ 0.4 turns per *μ*m, so formin will not generate supercoiling unless filaments become 6 *μ*m long, which we will not consider here. However, even the weaker twist *will* have an impact on the filopodial dynamics, and our goal is to determine how much this is.

#### 3.1.2 Buckling generated by polymerization without formin

Another way to get a nontrivial shape change in the central axis of the filopodium is polymerization. To illustrate this point, let us consider the five-filament bundle with all filaments having initial length *L* = 1 *μ*m. With the plus ends permanently crosslinked (imitating the filopodial tip complex), we allow the central filament to polymerize at rate 1 *μ*m/s until it reaches length *L* = 2 *μ*m. To make the dynamics reproducible (so buckling is not triggered by numerical roundoff), we make the central filament initially straight, but with tangent vector 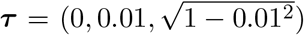 (all other filaments are exactly aligned with the *z* axis).

Fig 1C shows how the crosslinkers constrain the filament trajectory, inducing buckling. Initially, the central filament grows linearly, stretching the crosslinks. When the crosslinking force becomes too strong, the central filament buckles (at *t* = 0.2 s). After buckling, there is a balance of forces: the polymerization moves the filament end to the right, which takes the other filaments with it, bending the central axis of the bundle. When the bending forces of the other filaments become too large, these filaments stop bending, and the crosslinking forces hold the plus end of the central filament in place. As a result, polymerization is immediately pushed back, so that the filament grows outwards (towards the left). When allowed to equilibrate at a fixed length, the central filament finishes slightly above the other fibers, so that the downward crosslinking force balances the upward bending force on it (Fig. S1). In steady state, the average angle the filopodium makes with the initial axis stabilizes to roughly 15 degrees (Fig. S2).

#### 3.1.3 Active formin twist along with polymerization

To look at how the formin torque affects the bundle, we apply the torque of magnitude 0.1 pN·*μ*m to the growing filament (the torque becomes zero when the filament stops growing) and repeat the five-filament simulation. We observe dynamics which resemble those without formin, but with some small differences: the twisting appears to marginally accentuate trends we observed previously, with the mean filament curvature increase of ≈ 10% compared to the no-formin case (Figs. S3 and S4).

#### 3.1.4 Higher (supercoiled) twist density

Even though we will use the magnitude of formin torque equal to 0.1 pN·*μ*m, it is still instructive to look at what happens when we increase this torque beyond the threshold required for the supercoiling instability. Here we consider what happens to the same five-filament bundle when the torque is equal to 0.5 pN·*μ*m. Based on our linear stability analysis, we expect supercoiling when the filament length reaches ≈ 1.2 *μ*m.

The resulting deformations of the five-filament filopodium are shown in Fig. 1D. Based on prior results in this section, we might expect that increasing the twist angle to the supercoiling range will promote more bending of the actin shaft. What we find, however, is that a larger amount of twist (which is applied on 0 ≤ *t* ≤ 1, when the central filament is polymerizing) allows for more coiling of the *central* filament, which can then grow in length by increasing the length of coiling in the interior of the bundle, rather than buckling and pushing the peripheral filaments to the right. Thus, the results of this section suggest that a moderate amount of twisting leads to more curvature of the bundle, but twist which crosses the supercoiling threshold will actually cause the peripheral fibers in the bundle to bend less.

### 3.2 Variance in polymerization rates accelerates buckling, which is relieved by dynamic crosslinking

We now consider how the twist and curvature of the actin bundle evolve if all filaments in the bundle polymerize with different rates, which could happen if different filaments are associated with chemically different parts of the filopodial tip complex. For the purposes of illustration, we consider the same five-filament bundle as in Fig. 1, with permanent crosslinks at the filament plus ends. Each filament has a prescribed random growth rate which is drawn at *t* = 0 from a normal distribution with mean 0.5 *μ*m/s and varying standard deviation (we consider 0.2, 0.1, and 0.05 *μ*m/s), and then does not change in time. To prevent fibers from getting too long, we stop fibers from lengthening when the length gets longer than double the initial length of 1 *μ*m.

In Fig. 2A, we show the average statistics for filopodia growth with three different standard deviations in the growth rate (0.2, 0.1, and 0.05 *μ*m/s). To generate statistics, we run ten samples for each growth rate, then take the mean over those ten samples to get a mean statistic. The error bars are the standard error in the mean over five trials. Our statistics in Fig. 2A confirm the hunch that variance in the growth rates aids curvature. When the standard deviation is small (*σ* = 0.05 *μ*m/s), the fibers barely curve, and the average *z* coordinate (about 1.4 *μ*m) is essentially the same as if the fibers were all growing with rate 0.5 *μ*m/s (which would be 1.5 *μ*m). As we increase the standard deviation in the growth rate, we see more deviation from the straight shape, and higher average curvature.

**Figure 2:**
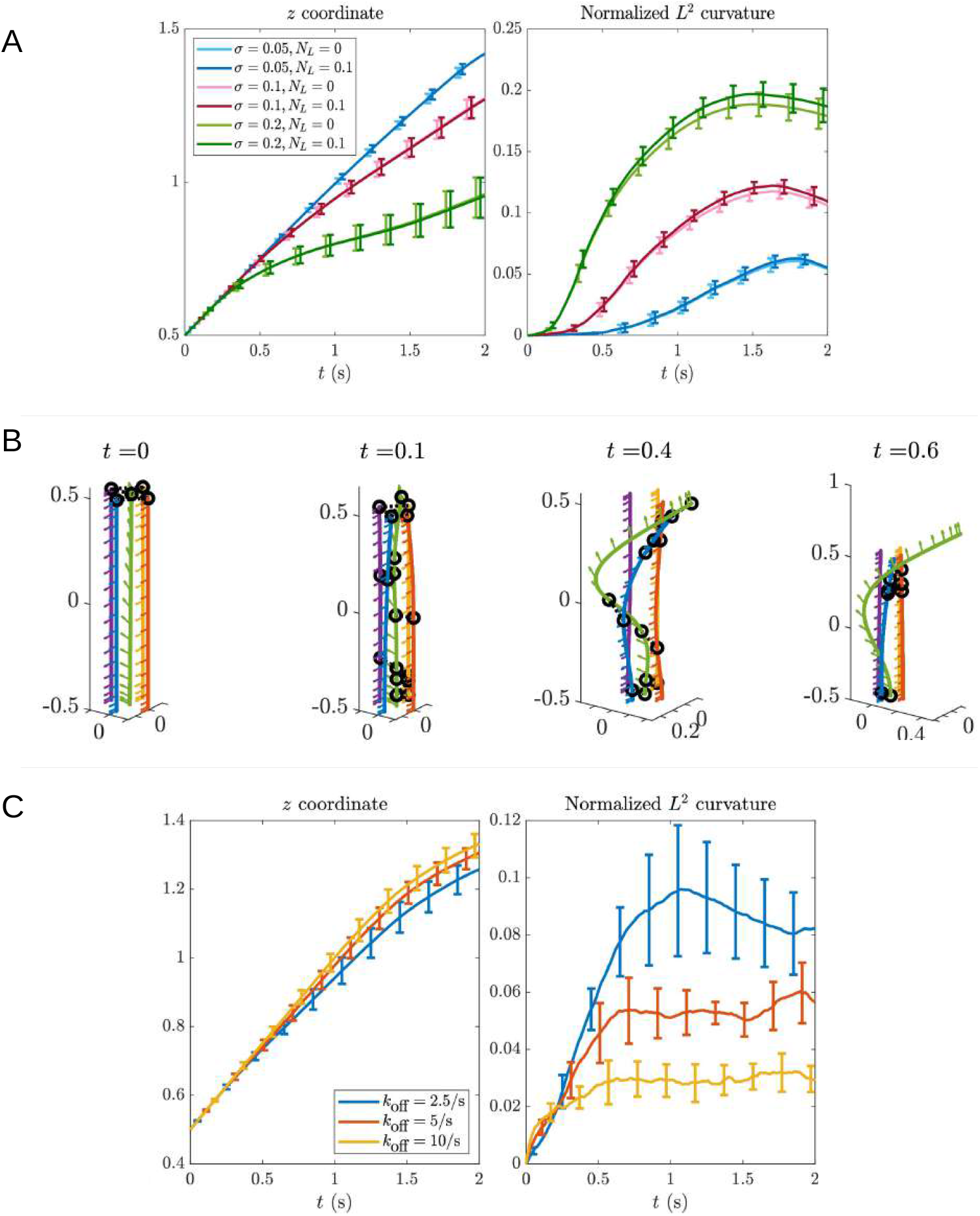
Lower variance in polymerization rates and dynamic crosslinking alleviate buckling. (A) Statistics for five-filament filopodial bundle with the tips of the outer filaments permanently crosslinked to the tip of the central filament. The growth rate of each filament is constant in time, but each filament’s growth rate is drawn from a normal distribution with mean 0.5 *μ*m/s and varying standard deviation (the values we use are *σ* = 0.2, 0.1, and 0.05 *μ*m/s). We plot the average *z* (in *μ*m) coordinate of the filament endpoints and the average filament curvature (normalized by the curvature of a circle with the same circumference as the filament length) as functions of time (in seconds). We are considering the mean over ten samples, with the error bars showing the standard error of the mean. The lighter colors are when no active torque is applied (*N*_*L*_ = 0), while the darker colors show the case when formin at the tip of each growing fiber generates a torque with value *N*_*L*_ = 0.1 pN·*μ*m. (B) Simulation of the five-filament bundle with polymerization of the central (green) filament and dynamic crosslinking. The polymerization is 1 *μ*m/s on 0 ≤ *t* ≤ 1 s, and dynamic crosslinks have on and off rates *k*_off_ = 5/s and *k*_on_ = 0.1/s. Arrows show the material frame vectors. Filaments are color-coded for visualization. (C) Statistics for five-filament filopodial bundle with transient crosslinkers. We fix *k*_on_*/k*_off_ = 0.02, so that the average number of links is roughly constant. Between the simulations, we vary parameter *k*_off_ (2.5/s – blue curves, 5/s – red curves, and 10/s – yellow curves). We plot the average *z* (in *μ*m) coordinate of the filament endpoints and the average filament curvature (normalized by the curvature of a circle with the same circumference as the filament length) as functions of time (in seconds). We are considering the mean over ten samples, with the error bars showing the standard error of the mean.

In Fig. 2A, we also consider the differences in the trajectories when we simulate without any formin torque (ligher colors), versus when the formin exerts a torque at the plus ends. In accordance with our observations in Section 3.1.3, we find that having a nonzero torque leads to fibers with higher average curvature. However, the effect is very slight.

#### 3.2.1 Dynamic crosslinking

We now switch on the crosslink dynamics to study how the bending behavior of the filopodium depends on the crosslinking turnover rate. We fix the crosslink dissociation rate at *k*_off_ = 5/s, then set the on rate *k*_on_ = 0.1/s so that there are about 10–15 links bound at any given time. (Technically, the on rate should have units of 1/(length*×*time) to be discretization independent. This is actually complex when the filament length changes in time, so we simply choose a value that gives a characteristic number of engaged crosslinks.)

We fix *k*_on_*/k*_off_ = 0.02, but vary the actual values of the rates, so that the average number of bound links stays roughly constant (at 8–10 links) while the residence time of the links changes. Fig. 2B shows an example when *k*_off_ = 5/s, where only the central filament polymerizes. We note how dynamic crosslinking alleviates the bending of the peripheral fibers – while the central fiber bends because of the initial constraint by crosslinking forces, the transient nature of the links prevents it from taking the peripheral filaments with it.

To quantify this effect, we repeat the test of the previous section (with a mean growth rate of 0.5 *μ*m/s and standard deviation of *σ* = 0.2 *μ*m/s), where we run 10 samples of the filopodial trajectories to get a mean, and repeat five times to generate error bars. Fig. 2C shows the mean bundle curvature and deviation from straight growth. As expected, links with longer residence time induce more bending of the filaments. Even at the longest residence time of 0.4 s (*k*_off_ = 2.5/s), we still observe curvature which is 2 to 3 times smaller than the value for permanent crosslinkers located at the filopodium tip (green lines in Fig. 2A). Thus, transient crosslinkers allow for more variable growth rates without strong bending of the central shaft. Note that there are two sources of bending in this case – one from the initial condition with the crosslinks at the tip, and another from the effective bending force arising when crosslinkers bind.

### 3.3 Motors can effectively braid filaments, aided by transient crosslinking

#### 3.3.1 Motors with longer range give more efficient braiding

We now add to the model of the previous section an external force density that describes the action of helical motors that act from the filopodial boundaries. It is important to remember that the motors can act only near the boundary – they cannot reach the central filament. We start by varying parameter *c*_*m*_, which is the length-wise fraction of the filopodium covered by the motors, with all other parameters fixed (this includes the condition that the motors only act along the outer 1*/*4 of the filopodial cylinder). The dynamics with *c*_*m*_ = 1 (motors covering the whole length) are shown in Fig. 3A, where we see that the fibers rapidly form a braided structure with multiple turns (more frames are shown in Fig. S5). After the plus end makes about two turns relative to the base, the braiding motion is arrested because fiber elasticity (which prevents the fibers from curving any more) and steric forces (which prevent the fibers from relaxing in the middle by crossing each other) balance the turning motion from the motors.

**Figure 3:**
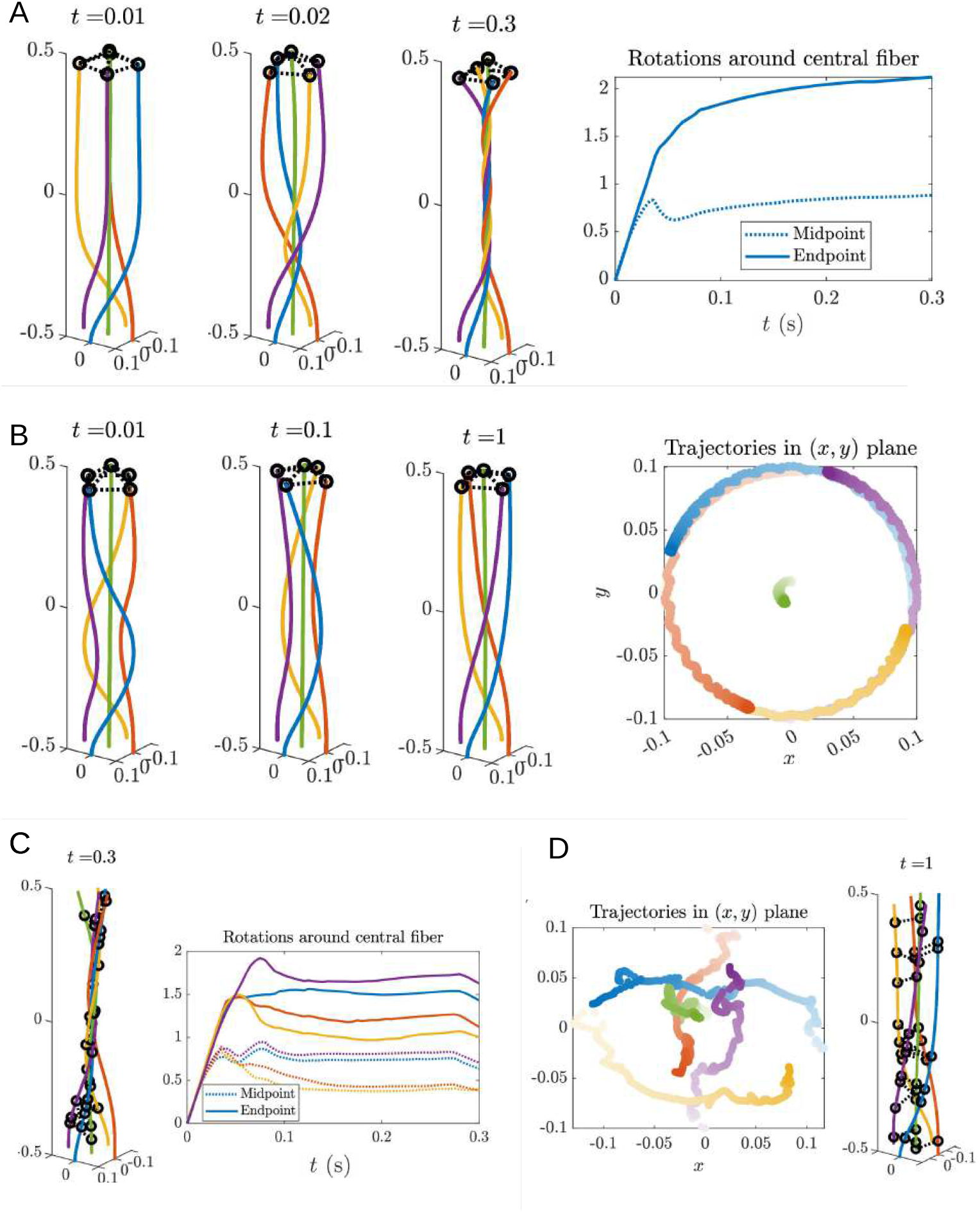
Motors acting on filopodium edges effectively braid the fibers, aided by permanent crosslinking in regions of motor activity. (A) Configurations of five-filament bundles when the motors act on the edges of the filopodium, and along the whole length of the filaments (*c*_*m*_ = 1). Crosslinks are dashed segments; rings are the endpoints of the crosslinks. Filaments are color-coded for visualization. Time is in seconds, distances are in *μ*m. The right-most image shows the number of rotations the peripheral fiber endpoint (solid line) or midpoint (dotted line) makes around the central fiber. More snapshots and statistics are shown in Figs. S5 and S6. (B) Configurations of five-filament bundle when the motors act only along *half* the length of the fiber (*c*_*m*_ = 1*/*2). The tips of the outer filaments are permanently crosslinked to the tip of the central filament. The right-most image shows a 2D projection of the fiber endpoints over time. The colors correspond to the fiber colors in the images at left, and darker dots show positions at later time points. More snapshots and statistics are shown in Figs. S7 and S8. (C) Ending configuration of five-filament bundle when motors act along the whole length of the filaments, (*c*_*m*_ = 1), and the crosslinks are transient. Initially, only the tips of the outer filaments are crosslinked to the tip of the central filament. The right panel shows the number of rotations each fiber’s endpoint (solid line of the same color) and midpoint (dotted line of the same color) make around the central filament. More simulation snapshots are shown in Fig. S9. (D) Ending configuration and projected endpoint trajectories of the five-filament bundle when motors act along *half* the length of the fiber (*c*_*m*_ = 1*/*2), where the crosslinks are transient. Initially, only the tips of the outer filaments are crosslinked to the tip of the central filament. More detailed dynamics are shown in Fig. S10.

**Figure 4:**
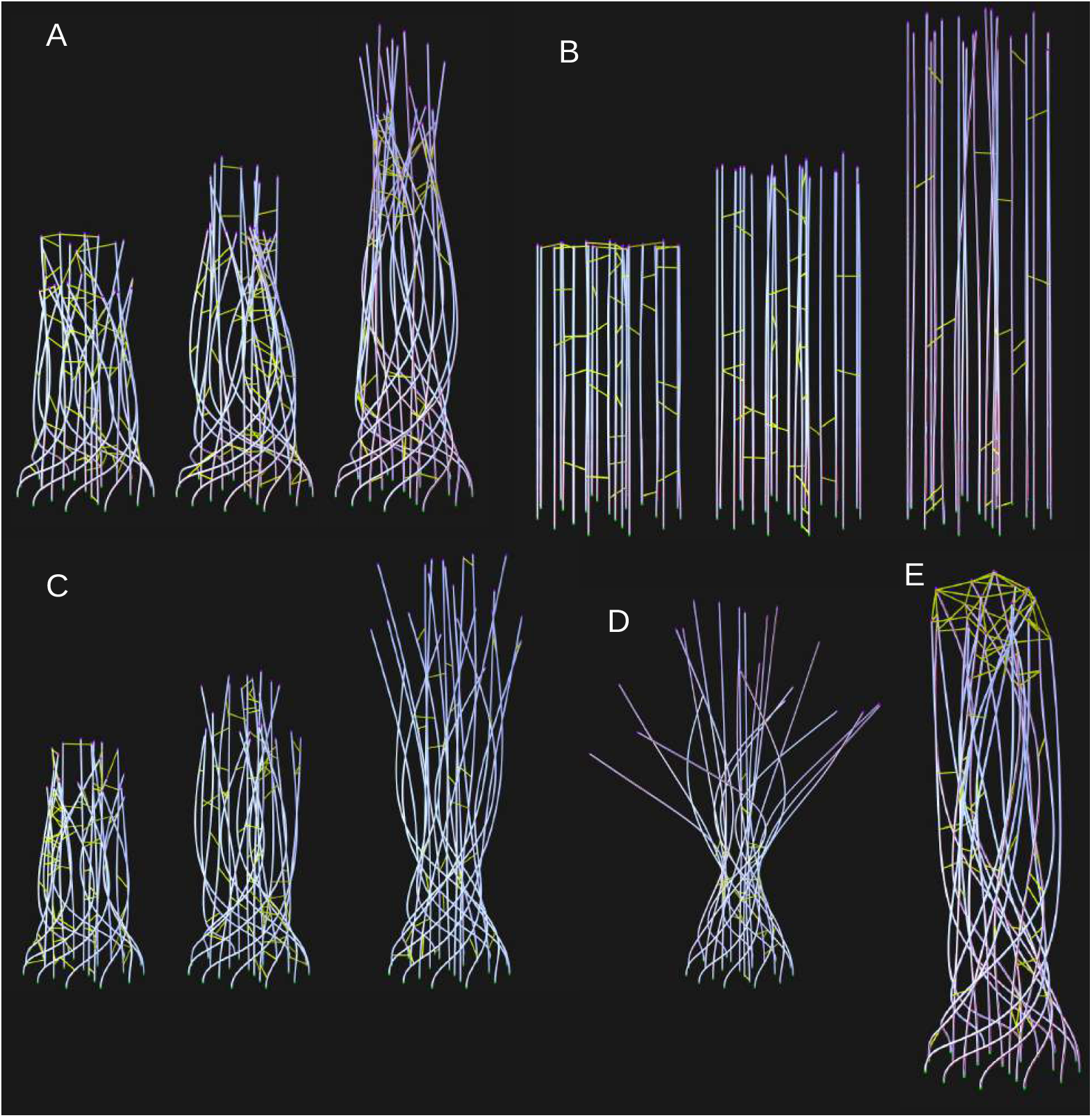
Snapshots of 22-filament bundles with and without motor and twisting activity, showing that motors and twist synergize to generate a compact bundle. Crosslinks are shown in yellow, filaments in white, and formins in magenta. The green dots show the clamped ends, which represent the branched actin network beneath the filopodium. (A) Time sequence of filopodium growth when there is both formin twisting and motors acting along the half length of the fiber; (*c*_*m*_ = 1*/*2). Snapshots are shown at *t* = 0.2, 0.8, and 2 s. See supplementary movie 1 for full sequence. (B) Time sequence of filopodium growth when there is only formin twisting, and no motor activity. Snapshots are shown at *t* = 0.2, 0.8, and 2 s. See supplementary movie 2 for full sequence. (C) Time sequence of filopodium growth when there is only motor activity, and no formin twisting. Snapshots are shown at *t* = 0.2, 0.8, and 2 s. See supplementary movie 3 for full sequence. (D) Snapshot from the end of a simulation (*t* = 2 s) when formins move in the opposite direction (CW, looking from the tip) from the motors (CCW, looking from the tip). See supplementary movie 4 for full sequence. (E) Snapshot from the end of a simulation (*t* = 2 s) when crosslinks at the tip are permanent (a sort of filopodial tip complex; in the simulations, permanent crosslinks connect the plus ends that are the nearest neighbors), while crosslinks below the tip are free to bind and unbind transiently. A less compact bundle with more bending and buckling is observed.

To get a better understanding of this process, we show the number of rotations the blue fiber (all other fibers are the same by symmetry) makes around the central green fiber as the last panel of Fig. 3A, for both the endpoint (plus end, solid line) and midpoint (dotted line). We see that the endpoint and midpoint first turn together, but around *t* ≈ 0.04 s the midpoint motion is arrested, and the endpoint motion is subsequently arrested around *t* ≈ 0.1 s. The rotation of the fibers combines with inextensibility to yield a downward motion of the *z* coordinate of the outer fibers. These in turn pull on the central fiber through the connected crosslinks, and the central fiber buckles slightly, curving and being pushed downward (see the full statistics in Fig. S6).

We now consider what happens when we change the motor forcing fraction to the lower 1*/*2 of the fiber. As shown in Fig. 3B, in the initial stages the action now happens in the middle of the fibers. These are pushed around in a circular trajectory, and the ends gradually follow via fiber elasticity. When the middle of the fibers is pushed into the middle of the filopodium, the motors can no longer access the fibers at all, and the trajectory slows down and stops. The statistics in Fig. S8 show that what results is oscillations, where the fibers go into the middle of the filopodium, losing access to the motors, then relax via fiber elasticity. These oscillations, which are also evident in the 2D projection of the fiber endpoints in Fig. 3B, occur when elastic relaxation takes a piece of the fiber into the motor zone, which forces it again back to the middle. Fig. S8 shows that this process essentially repeats over *t* ≥ 0.5 s, yielding little changes in the trajectory. We notice also that the motors do not provide enough force to buckle the central fiber at all (Figs. S7 and S8).

#### 3.2.2 Transient crosslinking extends the reach of motors

We now look at what happens when the crosslinkers binding the filaments are transient. We will in particular consider the same five-filament set-up, initially with *c*_*m*_ = 1, so that the motors are acting their strongest (along the length of the whole fiber). We then introduce transient linking with constants *k*_off_ = 10/s (we choose the fast rate because the braiding process for *c*_*m*_ = 1 can be quite fast – timescale 0.1 s; see Fig. 3A – and we want the links to be transient on those timescales).

Compared to the case with permanent crosslinkers (Fig. 2A), we see that the fibers are not as twisted, as the peripheral fibers in the braid turn 1–1.5 times around the central one (Fig. 3C, see also Fig. S9 for the full time sequence). The reason for this is that there is less force from the motors. When the links near the top of the fibers unbind, those parts of the fiber tend to relax into the center of the filopodium. This takes them out of reach of the motors, so that by *t* = 0.3 s the top parts of the fiber move around via crosslinking action, and not motors. This renders a dynamic steady state.

Given this observation, we might expect transient crosslinking to hinder braiding when *c*_*m*_ = 1*/*2 also. However, because the motors do not act along the top of the fiber when *c*_*m*_ = 1*/*2, unbinding of the links has no effect there. Indeed, as shown in Fig. 3D (full time sequence in Fig. S10), the configuration of the filopodium with *transient* linkers is roughly the same as with permanent linkers, and the 2D projection of the endpoints shows that the number of turns the peripheral fibers make is roughly the same. Thus, for a filopodium of this size, transient linking makes little difference in the overall dynamics.

### 3.4 Formins and motors synergize to twist and compress large filopodia

After building intuition with small bundles, we turn to a larger bundle with 22 fibers (characteristic for filopodia) arranged into four rings (1, 3, 6, and 12 filaments). The rings are spaced 100 nm (the crosslinker rest length) apart, and fibers are positioned with uniform spacing around each ring. Starting with this initial geometry, we repeat the simulations of the previous section, with the following modifications: The average growth rate of each filament is the same, 0.5 *μ*m/s, but the growth rate of each filament is a random variable which varies in time according to Eq. (1). In practice, this yields a normal distribution of growth lengths at any time with standard deviation 0.05 *μ*m/s. In these simulations, the motors are always distributed in the bottom half of the filopodium (*c*_*m*_ = 1*/*2). The crosslinkers are dynamic with on rate *k*_on_ = 0.2/s and off rate *k*_off_ = 10/s.

What we are interested in is how the motor activity and twisting activity at the tip affect the overall chirality of the bundle. By twisting activity at the tip, we mean the torque *N*_*L*_ = 0.1 pN· *μ*m that the formin exerts on the fiber tip. Thus, we will consider three sets of simulations: one with twisting and no motor activity (shown in Fig. 4B) one with motor activity and no twisting (Fig. 4C), and one with both twisting and motor activity (Fig. 4A). These plots show three time points (*t* = 0.2, 0.8, and 2 s); for more time points see the supplementary movies.

Without motors, the twisting of the filaments at the tips makes little difference (Fig. 4B). We observe filaments which basically grow straight outward, segregating from time to time into mini-bundles as dictated by the crosslinkers. The bundle has no discernible chirality. When we add motors, but remove twist, we see a braid coming together, but the packing appears fairly loose (Fig. 4C). By contrast, when we add formin twisting to the motor activity, we see a bundle which is significantly tighter and with filaments that are wound more around the central fiber (Fig. 4A). To confirm that the qualitative packing in the bundles is due to the physics of the motor-formin interaction and not random chance, we perform ten simulations for each of the three parameter sets and compute a mean and standard deviation for each of our usual statistics. Unlike in simulations with five fibers, here we expect the location of a fiber (in terms of which ring it sits in) to be important for determining the statistics of its motion (since, e.g., motors can only act along the outer 25% of the bundle, which means they can only act along the outer circle of fibers initially). In Figs. S11–S12, we therefore segregate our statistics by ring or circle, showing the central fiber in blue, the smaller circle of 3 fibers in red, the middle circle of 6 fibers in yellow, and the largest outer circle of 12 fibers in purple.

Beginning with the mean curvature (Fig. S11), we see a marked contrast between simulations without and with motors. Without motors, the filaments barely curve at all, and more curvature emerges the closer the fibers are to the bundle center, where they presumably can access more of the peripheral filaments via crosslinking. With motors, the largest curvatures happen at the periphery, since the outer circle of filaments (in purple) are twisted by the motors while their ends remain clamped. When adding twist, there is a small (∼ 10%), but significant, increase in the curvature of the outer circle of filaments, consistent with the fibers having to bend more to align themselves more tightly at the filopodium tip.

The number of rotations around the central fiber statistic also shows a contrast between simulations with and without motors, and a smaller but still significant difference in simulations with motors but with and without twist. As expected, we observe little twist without motors (Fig. S12, left). When we add motors, the twist angles grow as we move outward from the inside to the periphery of the filopodium (Fig. S12, middle). The bundles with formin twist also appear to be more successful at rotating the inner rings of filaments (compare yellow and red lines in the right two panels of Fig. S12) around the central filament: when the outer filaments are more tightly aligned, more crosslinks can form at the top of the bundle, which allows filaments in the interior to be transported with the filaments at the periphery (since only the ones at the periphery are acted on by the motors). This provides another example of why formin twist is vital for filopodium stability.

#### 3.4.1 The twisting formin pushes the filaments into the center of the filopodium

To verify that the formin twist causes inward motion of the peripheral fibers, we study the final position (at *t* = 2 s) of the peripheral filament plus ends across our ten simulations. In Fig. 5A we plot the endpoints of the outer 12 fibers in the filopodium, projected onto the (*x, y*) plane. Since there are ten trials, each color/marker type has a total of 120 observations. In addition to plotting simulations with CCW formin twist and no motors (blue circles) and motors and no twist (black diamonds), we plot simulations with motor activity and twisting. The red squares show when the formin torque is applied in the same CCW direction as the motors, while the green triangles show the endpoints when the torque is applied in the *opposite* CW direction (Fig. 4D gives an impression of the end position of sample simulations). The significant change from a compact set of the plus ends’ ending positions (when the torque is in the same direction) to a very large outward splaying of the plus ends’ ending positions (when the torque is in the opposite direction), shows how formin twist in the same (CCW) direction as motor spin is vital to maintain a tight filopodium shape. Indeed, the filopodium with motors and twist is actually tighter than without motors!

**Figure 5:**
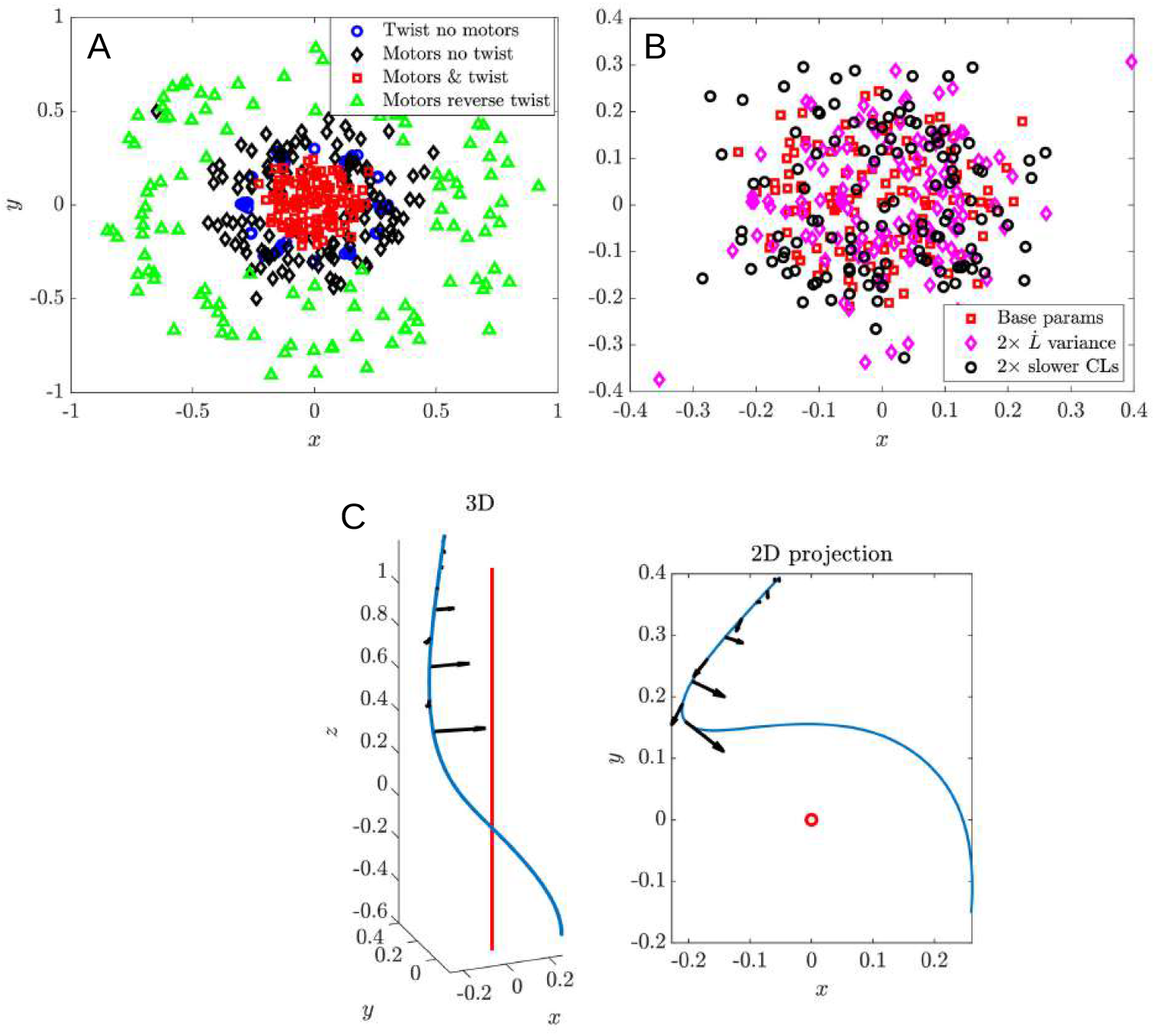
Compaction is driven by twist forces (activated by bending), and is most efficient when the crosslinks are more transient and the filament growth rate is roughly constant. (A) The positions of the outer 12 filaments’ tips at *t* = 2 s, projected onto the (*x, y*) plane for the three simulation conditions. Blue circles correspond to twist without motors, black diamonds correspond to motors without twist, and red squares correspond to motors with twist. The green triangles correspond to what happens when we reverse the direction of formin twisting. Simulations were repeated 10 times for each parameter set. (B) The same projection as in (A), but with different parameter sets. Red squares again show the base parameters, while pink diamonds show simulations with a twice larger variance in the polymerization rate 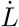, and black circles show simulations with twice slower on and off rates for the CLs. (C) Shape of an outer filament (in 3D and projection onto the (*x, y*) plane) in the 22-filament filopodium as computed when the motors apply force to the lower half of the bundle, but without the formin twist. Then, we apply the formin twist to such filament, and show the resulting forces (arrows) generated by the twist on the upper region of the filament. The red line (circle in the 2D projection) shows the central (*z*) axis of the filopodium.

To understand this effect, we isolate a filament *from a simulation without twist* and compute the twist force:

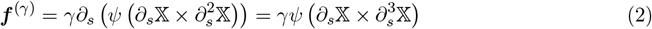

when the twist angle *ψ* = *N*_*L*_*/γ* is in its constant steady state (this is how the second equality follows from the first). Here *γ* is the torsional rigidity of actin, *N*_*L*_ is the formin torque, and X is the 3D curve of the filament centerline (see the SM for details). The fiber and the twist force density on its top half (where motors do not act) are shown in Fig. 5C. There, we see that the twist force is largely directed towards the center of the filopodium, as is qualitatively clear from the large bundle simulations. Eq. (2) also shows why the twist force is only significant when motors bend the fibers: the force density is only nonzero when 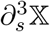 is nonzero! Thus, when fibers are straight, as is the case when motors are absent, the twist force is zero, and there is no way for the twist to feed back onto the fiber shape. It is necessary for the motors to first bend the fibers, and then formin twist can affect the shape. Thus, the motors and formins synergize to make the bundle more compact.

#### 3.4.2 Transient crosslinking and constant polymerization rates aid filopodium compaction

We now vary the rate of crosslinking turnover and filament polymerization. We consider a longer crosslinking residence time, decreasing the rates of crosslink attachment and detachment by a factor of two, and a larger, by a factor of two, standard deviation in the filament polymerization rate. In Fig. 5B, we plot the plus ends of the peripheral circle of fibers as compared to our base simulations. We see that adding variance in the polymerization rate can induce outliers in the final plus end position, and that a longer crosslink residence time induces a distribution of plus ends that is wider than the original parameters. This tells us that dynamic crosslinks connecting the inner filaments to the outer ones also play a role in drawing the bundle inward, and that a more uniform polymerization rate helps keep a compact bundle.

#### 3.4.3 Influence of the filopodial tip complex

As an example of what can happen when the plus ends are associated with formins, which in turn are associated with a solid filopodial tip complex, we consider a simulation where the crosslinks at the fiber plus ends are permanent, mimicking the filopodial tip complex, while crosslinks connecting the filaments’ sides can form and break dynamically. Fig. 4E shows the resulting filopodium. Comparing to Fig. 4A, we observe significantly more bending of the interior fibers and a much wider bundle, as the filopodial tip complex provides an additional constraint preventing the filaments from being drawn into the middle. In addition, the filaments appear more buckled because of the variance in the polymerization rate, and there is a direction of bending to the filopodium, similar to what we saw earlier in five-filament bundles with permanent crosslinks.

## 4. Discussion

We found that the spinning action by helical motors effectively “braids” the actin bundle, compacting it, generating buckling, and enhancing the crosslinking. Stochastic fluctuations of actin polymerization rates also contributed to filament buckling and bending of the bundle. Faster turnover of transient crosslinks attenuated the buckling but could enhance coiling and compaction of the bundle. Interestingly, formin-generated twisting alone was much less effective than myosin-generated spinning in inducing filopodial braiding and compaction, but formin action helped the motors to compact the bundle once braided.

Our model prediction that myosin action, as opposed to formin activity, is the key for coiling the filopodium, is in agreement with the observation that the molecular activities of myosin V and X are involved in rotation of filopodia whereas the formin mDia1 does not play a role in the observed rotations (Leijnse et al., 2022). We note though that formins may not rotate or twist filaments (Kovar and Pollard, 2004), which could be an alternative explanation of why perturbations of formin reported in (Leijnse et al., 2022) did not inhibit the chiral filopodial structure. Another observation from our simulations – that buckling of several filaments that grow faster than others can create an actin loop and respective bulge at the filopodial tip – could be relevant to similar experimentally-observed structures (Li et al., 2023). The mechanisms we discussed here could also be relevant to processes that trigger chiral tilting of actomyosin fibers in adherent cells (Tee et al., 2015). Some of these mechanisms are very similar to microtubule/kinesin interactions generating chirality of the mitotic spindle (Novak et al., 2018).

One of the model predictions – that faster crosslink turnover enhances coiling and compaction in the filopodium – is nontrivial and probably could be tested in future experiments. Note though that here we tested crosslinking with very rapid kinetics on the scale of several cycles of binding/unbinding per second, which is characteristic of the ubiqitous crosslinker *α*-actinin (Wachs-stock et al., 1994). The principal crosslinker in filopodia, fascin, has much slower kinetics with the turnover cycle on the scale of 10 seconds (Aratyn et al., 2007). Another interesting prediction for the future is that the filopodial bundle chirality is the same as that of the helical myosin motors: if the motors spin filaments CW (CCW), then the bundle will coil CW (CCW), respectively. (An open question is what happens if there is a mixture of motors with opposite helicity.) The predicted effect of formin depends on the relative helicity of the motors and formin: if formin-generated torque rotates filaments in the same direction as the motors, then the bundle will be tighter; if the formin torque is opposite to the motor spinning action, then the bundle will be more disheveled.

What could be the possible roles of the spinning and twisting actions of the myosins and formins, respectively, and of the resulting coiling and buckling of the filopodial bundles? One very suggestive role, based on the simulations, is the compaction of the bundle, which likely means greater mechanical stability. Mechanical stability of the bundle is, in fact, affected by coiling even without compaction (Daniels and Turner, 2013). Another possibility is that the spinning action could result in rotation of the filopodia around axes not coinciding with bundle axes, as was observed in (Tamada et al., 2010), which could allow filopodia scanning over a greater area around the cell edge. Some filopodia fold back into the cell leading edge and contribute to construction of contractile bundles in the lamella (Nemethova et al., 2008), and in that way pre-established filopodial chirality could trigger larger scale chirality of the cell cortex by interaction with other cytoskeletal structures of the cortex. Coiling and buckling in the filopodial bundle can induce pulling and traction at the tip (Leijnse et al., 2015). Last, but not least, straightening of the coiled actin deformation could generate protrusion and force, as in the dramatic acrosome reaction of *Limulus polyphemus* sperm (Shin et al., 2003).

The mechanism of chirality emergence that we explored is but one possibility, and there is likely a diverse inventory of chirality propagation mechanisms, even for just actin bundles. An unknown mechanism, not relying on either myosins or formins, is responsible for a chiral rotation of *Listeria* propelled by a polarized actin tail (Robbins and Theriot, 2003). Turning of long filopodia to the left (Tamada et al., 2010; Li et al., 2023) is not explained by the mechanisms that we considered. General thermodynamic arguments demonstrated that when chiral filaments are bundled, a macroscopic twist is generated in the bundle (Grason and Bruinsma, 2007). It was shown theoretically that stereospecific crosslinking in parallel bundles of helical filaments, such that possible points of crosslink binding to actin filaments are located along helical patches on the filaments and are not completely flexible, generates intrinsic torques, of the type that tend to wind the bundle superhelically about its central axis (Heussinger and Grason, 2011; Grason, 2015). Recently, a detailed simulation with monomer-scale resolution confirmed that torsional compliance in a finite-width filament can induce chirality in the structure of a crosslinked bundle (Floyd et al., 2022). One of the interesting consequences of such a mechanism is that bundles that are too thick become destabilized due to the energetic trade off between filament twisting and crosslinker binding within a bundle. In fact, it was experimentally observed that small rigid actin-binding proteins change the twist of filaments in a concentration-dependent manner, resulting in small, well defined bundle thickness up to 20 filaments (Claessens et al., 2008), which, accidentally or not, is on the order of the filament number in filopodium. Finally, actin bundling by counterions is also predicted to generate chirality (Mohammadinejad et al., 2012).

Our model, though overly simplified, produces a rich variety of filopodial deformations, so we expect that adding factors that we ignored will predict even more nontrivial possible behaviors. These factors include, but are not limited to: 1) complex structure of the filopodial bundle, in which the filaments do not grow continuously from the base to the tip, but rather arrays of distinct filaments overlap along the bundle length (Medalia et al., 2007); 2) mechanical coupling of twisting and bending, even for a single filament (De La Cruz et al., 2010); 3) complexity of the connections of the filopodial filaments to the lamellar actin network (Evans et al., 1997; Svitkina et al., 2003); 4) complex structure and mechanics of the filopodial protein tip complex (Cheng and Mullins, 2020); 5) formin sensing both force and torque during actin filament polymerization (Yu et al., 2017); 6) mechanical interactions of the actin bundle with a membrane enveloping the bundle, which would affect both buckling and coiling (Pronk et al., 2008; Daniels and Turner, 2013). Inclusion of some of these factors into the model and scaling up the simulations will allow us to address the open question about how chirality propagates to the cellular (Tee et al., 2015; Zaatri et al., 2021) and multicellular scales (Yashunsky et al., 2022; Tee et al., 2023).

## Acknowledgements

This work was supported by the National Science Foundation through Research Training Group in Modeling and Simulation under award RTG/DMS-1646339 and through the Division of Mathematical Sciences awards DMS-2052515 and DMS-1953430. Ondrej Maxian was supported by the NSF via GRFP/DGE-1342536. We thank A. Bershadsky for help with literature and fruitful discussions.

## Competing interests

The authors have no competing interests to declare.

## Supplemetal Material for

### 1. Filaments as inextensible bent and twisted rods

We first summarize our previous work on twisting filaments, highlighting changes since the publication of [16], on which most of our numerical method is based. We let 𝕏(*s*) be a parameterization of the filament centerline, and introduce an operator **𝒦** [𝕏(·)] which parameterizes the space of inextensible filament motions. The translational velocity can be written in terms of such motions as

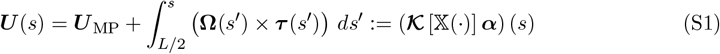

where ***U*** _MP_ is the velocity of the filament midpoint and ***α*** = (**Ω, *U*** _MP_) represents the translational degrees of freedom. In continuum, the dynamics of the centerline are then governed by the saddlepoint system

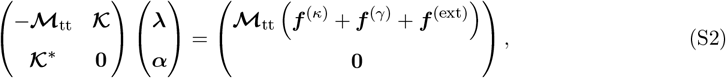

where

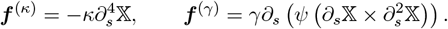

The first equation says that the total velocity is obtained by multiplying the total force density, which is composed of the constraint force ***λ***, the bending force ***f*** ^(*κ*)^, the twisting force ***f*** ^(*γ*)^, and any external forcing ***f*** ^(ext)^, by the mobility **ℳ**_tt_. The second equation, **𝒦**^*^***λ*** = **0** enforces the principle of virtual work, which says that the constraint forces ***λ*** perform no work with respect to all inextensible motions. As discussed in [16, Sec. III(A)], this constraint is equivalent to setting ***λ*** = *∂*_*s*_ (*T* ***τ***), for some line tension *T* (*s*) [23].

The twist density on the filament centerline evolves via an auxiliary set of equations. The total parallel torque is given by

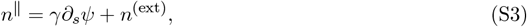

where the first term is the parallel torque due to twisting, and the second one represents externally-applied torques (e.g., from motors or crosslinkers). The twist-density *ψ* then evolves according to the equation

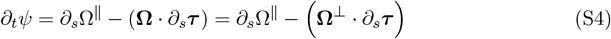

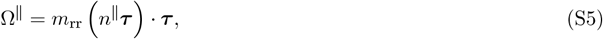

is the parallel rate of rotation, the perpendicular rate **Ω** is obtained from the solution of (S2), and *m*_rr_ represents the mobility (inverse of drag) relationship between (parallel) torque and (parallel) angular velocity.

Our simulations use a clamped end at the filopodium base (*s* = 0),

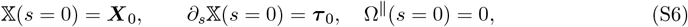

where ***X***_0_ and ***τ***_0_ are the fixed endpoint position and tangent vector, respectively. At the clamped end, we assume that the parallel angular velocity is zero,

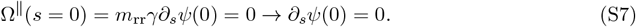

The *s* = *L* is more delicate. There, we assume that a clamped formin exerts a fixed *torque N*_*L*_ on the fiber,

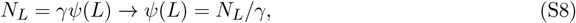

while the position of the fiber end is free,

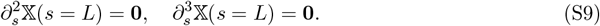

In this work, we are interested in capturing the qualitative behavior of the filaments, but not necessarily the quantitative hydrodynamics. Because of that, we will neglect all hydrodynamic interactions between filaments (and between far-away parts of a single filament), and any coupling between rotation and translation (i.e., we assume that the torque does not induce a translational velocity; see [16] for more details). For the purposes of this study, we only want to capture the qualitative separation of scales between bending and twisting. Because of this, we use the local drag mobilities [20]

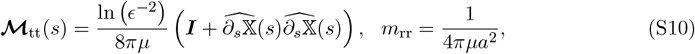

where *a* is the filament radius and *ϵ*(*t*) = *a/L*(*t*) is the filament aspect ratio. Here the translational mobility **ℳ**_tt_ is a 3 *×* 3 matrix on each filament point, which expresses the fact that the resistance in the tangential direction is half that in the perpendicular direction. The rotational mobility *m*_rr_ is a scalar that relates parallel torque *n*^∥^ to rotational velocity Ω^∥^. These mobilities are the minimal number of quantities necessary to capture the separation of scales (twist equilibrates **𝒪**(*ϵ*^−2^)-fold faster than force).

Because all quantities are ultimately discrete in simulations, we discretize the continuum curve 𝕏(*s*) using a set of *N*_*x*_ Chebyshev nodes, which we denote by ***X***. These nodes define an interpolating function, which can be used at any point to obtain the position 𝕏(*s*) (see [15, Sec. 1] or [11, Sec. 6.1] for an explanation of our discretization). Once we have a spatial discretization, the saddle point system (S2) becomes a discrete matrix equation that we solve at each time step. To make our time-marching scheme stable for larger time steps, we treat the bending force implicitly; see [11, Sec. 6.4] for more details.

### 2. Crosslinking

To compute crosslinking forces between pairs of filaments, we use the methodology described in [15, Sec. 6] to transform forces on a grid of uniformly spaced points along the filaments to forces on the Chebyshev grid. The forces can then be converted to the force density ***f*** ^(CL)^ by multiplying by an inverse weights matrix.

For the uniform points, we resample the filament 𝕏(*s*) at *N*_*u*_ uniformly-spaced locations, obtaining a vector 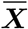 of sites where crosslinks are bound. The relationship between the two is given by

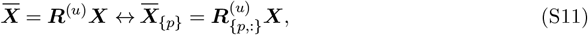

where ***R***^(*u*)^ is the resampling matrix and 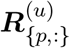 is its *p*th row.

Denoting the uniform points connected by the crosslink with 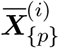 and 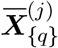, we have the displacement

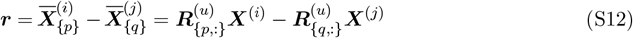

with *r* = ∥***r***∥. Then, let us postulate the crosslink energy and force,

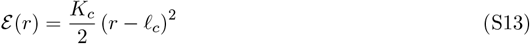

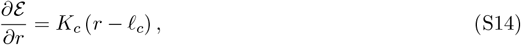

respectively, between the two uniform points, where *K*_*c*_ is the spring constant for the crosslink (units force/length) and *ℓ*_*c*_ is the rest length. The corresponding force on filaments *i* and *j* can then be obtained by differentiating the energy

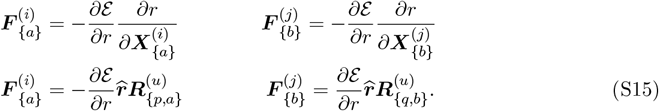

Note that in the model the crosslinks are linear springs that connect to the filaments by flexible joints, so possible bending forces are ignored. Also, the crosslinks bind, effectively, to the 1D contour of a filament, so potential effects from the specific distribution of binding sites along the helical grooves of the filaments are also ignored.

### 3. Steric forces

We compute steric interaction forces using the method described in [12, Sec. 9.2]. In brief, the steric interaction energy between filaments *i* and *j* can be written as

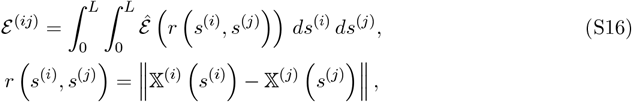

where 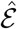 is the potential density function

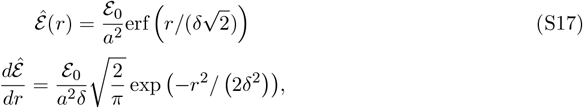

where the Gaussian is truncated at *r*_max_ = 4*δ*, and **ℰ**_0_ = 4*k*_*B*_*T* [22] is the magnitude of the steric force, which leaves *δ* = *a* is the parameter that controls the Gaussian decay.

To determine the forces, we compute the double integral (S16) for energy via upsampling the position to a high-resolution Chebyshev grid, then differentiate to get force. Similar to the cross-linking force, we denote the upsampled points via 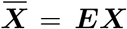. The double integral can then be evaluated and differentiated via

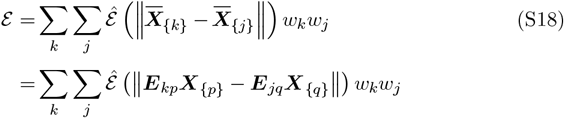

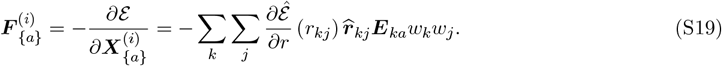

The last equation gives the force at Chebyshev node *a* on filament *i*, and is a function of the integration weights *w*_*k*_ and *w*_*j*_ of points *k* and *j* on the upsampled grid. In this work, we use *N*_*u*_ = 100 upsampling points to compute the steric forces on the filaments, and the points we choose are at arclength coordinates *s* = 0, Δ*s*_*u*_, …, *L*, where Δ*s*_*u*_ = *L/*(*N*_*u*_ − 1) is the spacing. The corresponding weights are *w* = Δ*s*_*u*_*/*2, Δ*s*_*u*_, … Δ*s*_*u*_, Δ*s*_*u*_*/*2, so that the first and last point have a weight of 1/2, in accordance with the trapezoid rule. Note that the sum (S19) can be computed efficiently via a neighbor search (rangesearch in Matlab).

### 4. Motor forces

We initially position the filaments around a circle of radius *R*, with all tangent vectors aligned with the *z* axis. To compute motor forces at a given time, we transform the coordinates of filament point (*x*_*p*_, *y*_*p*_, *z*_*p*_) to obtain an angle on the unit circle via Θ = arctan (*y*_*p*_*/x*_*p*_). The force density applied by the motor at that point is then ***f***_*p*_ = *f*_0_ (− sin Θ, cos Θ, 0), which keeps the fiber moving around the unit circle and has magnitude *f*_0_. These forces and torques are applied over a fraction of the fiber *c*_*m*_, i.e., over an interval [0, *c*_*m*_*L*] in filament arclength.

### 5. Formin-generated twist and supercoiling

This section provides a linear stability analysis of the behavior that leads to actin supercoiling. Because twist equilibrates much faster than bending, we can say that the twist evolution equation (S4) is in a steady state, so that it obeys the two-point boundary value problem

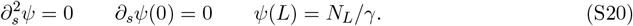

This boundary value problem has the obvious solution

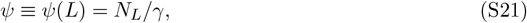

for the twist profile. The constant twist (S21) implies that the angle of rotation of the fiber *θ* = *ψ*(*s*) *ds* = *ψs*, is linear along the fiber.

Armed with this, we can perform a stability analysis on the fiber evolution equation. To do this, we linearize the evolution equation (S2) around a straight fiber 𝕏_0_(*s*) = (0, 0, *s*), then substitute the constant *ψ* from (S21). Setting 𝕏 = 𝕏_0_ + *δ𝕏*^⊥^, where *δ𝕏* is a *perpendicular* perturbation, we get the linearized dynamics (to first order in *δ* in the perpendicular direction)

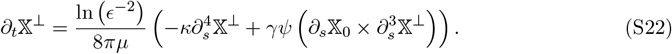

We now follow the technique of [27] by mapping 𝕏^⊥^ = (*X, Y*, ∼) to the complex plane, so that the complex number *h* = *X* + *iY* represents the degrees of freedom, and 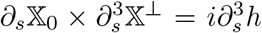. The transformed evolution equation (S22) becomes

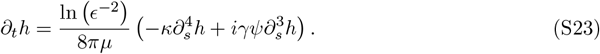

For linear stability analysis, we set *h*(*s, t*) = *e*^*ωt*^*ξ*(*s*), which gives the two point eigenvalue problem

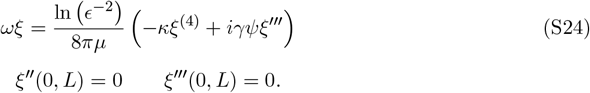

Our goal here is to find *ψ* such that the perturbation is linearly unstable. Similar to [27], we expect that, at the critical *ψ* = *ψ*_crit_, the twisting torque ∼ *γψ* will be equal to the bending torque ∼ *κ/L*. As such, we look for a critical value of *ψ*_crit_ = *cκ/*(*Lγ*) such that there exists an eigenvalue *ω* in (S24) with positive real part. This can be done numerically using Chebfun [19, 3], an open-source software for numerical computation with functions, which gives

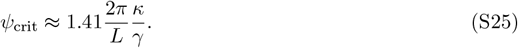

Converting this to a number of turns, we find that *N*_t,crit_ = *Lψ*_crit_*/*(2*π*) = 1.41(*κ/γ*). For actin filaments, *κ* ≈ 17 pN·*μ*m^2^ [5], while *γ* ≈ 10 pN·*μ*m^2^ [2]. This gives a critical number of turns *N*_t,crit_ = 2.4 for actin filaments of given length. If a filament is twisted less than that, it remains straight; if it is twisted more, it supercoils.

A similar stability analysis to the one here was also conducted for a clamped filament being spun at one end [27], and a twisted closed loop [26, 21].

### 6. Numerical method for polymerization

To model linear polymerization of a filament 𝕏(*s*) at the *s* = *L* end, we use the following algorithm:

1. Sample the Chebyshev interpolants 𝕏(*s* = *L*) and *∂*_*s𝕏*_(*s* = *L*) to obtain the position and tangent vector at *s* = *L*. Let ***X***_*L*_ denote the position and ***τ*** _*L*_ denote the normalized tangent vector.
2. Add an additional point to the matrix of Chebyshev nodes,

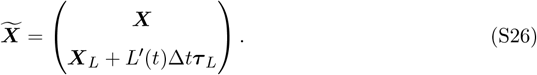
3. Reparameterize the filament by its new length. That is, the new set of evaluation points in *s* is given by 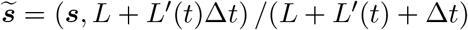 and goes from zero to one.
4. Let ***R*** be the matrix that evaluates the Chebyshev interpolant defined by the arclength points ***s*** at ***s***. Then set 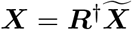 to obtain the positions at the Chebyshev nodes at the next time step. This resamples the filament at the nodes ***s***, but in the new parameterization.
5. The parameter *s* is no longer an arclength parameter, but is the distance in *reference arclength* coordinates. The new arclength parameter is given by

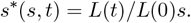

Thus, at the next time step, we can solve all the equations in coordinates of Section 1 via the transformation *∂/∂s*^*^ = (*L*_0_*/L*(*t*)) *∂/∂s*, i.e., by scaling all differentiation and integration operations appropriately.

We note that a similar reparameterization algorithm was employed in [18, Sec. 2.5] to model polymerization. However, in that work, the authors used the chain rule to write

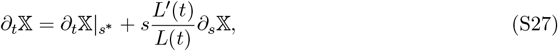

so that the first term can be obtained using an inextensible algorithm, and the second term added on afterwards. We found this formulation to be highly unstable numerically because each point locally extends in a direction according to its tangent vector. Because of this, numerical instabilities in tangent vector directions tend to be amplified. Our approach is more physical, because it models the addition of monomers at the *s* = *L* end, which happens along the direction corresponding to the last tangent vector. As (S27) suggests, we also first solve the equations of Section 1 to evolve the filament positions (inextensible in the *s*^*^ coordinate frame), then add the polymerization at the end of the time step.

### 7. Dynamic crosslinking

To implement dynamic crosslinking, we use a simplified version of the algorithm described in [14, 13]. Each filament is discretized into *N*_*u*_ = 41 binding sites. We assume that crosslinkers have rest length *ℓ*_*c*_, and can stretch at most by a length 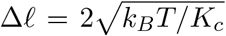. Thus, at each time step, we find all pairs of uniform sites on distinct fibers separated by a distance between *ℓ*_*c*_ − Δ*ℓ* and *ℓ*_*c*_ + Δ*ℓ*, and assign to each pair a binding rate 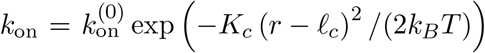 . We also assign each already bound link an unbinding rate of *k*_off_. This gives a set of reactions and rates, which we simulate at each time step using Gillespie’s next reaction method [1, 4]. To simplify the algorithm, we assume that unbound links will not rebind in a single time step, and that bound links will not unbind in a single time step. Our previous work had a more detailed model where each end of a crosslinker is distinct; here we simply treat the crosslink as one entity that is bound or unbound.

### 8. Parameters

Throughout this study, we use the parameters in Table 1.

**Table 1:**
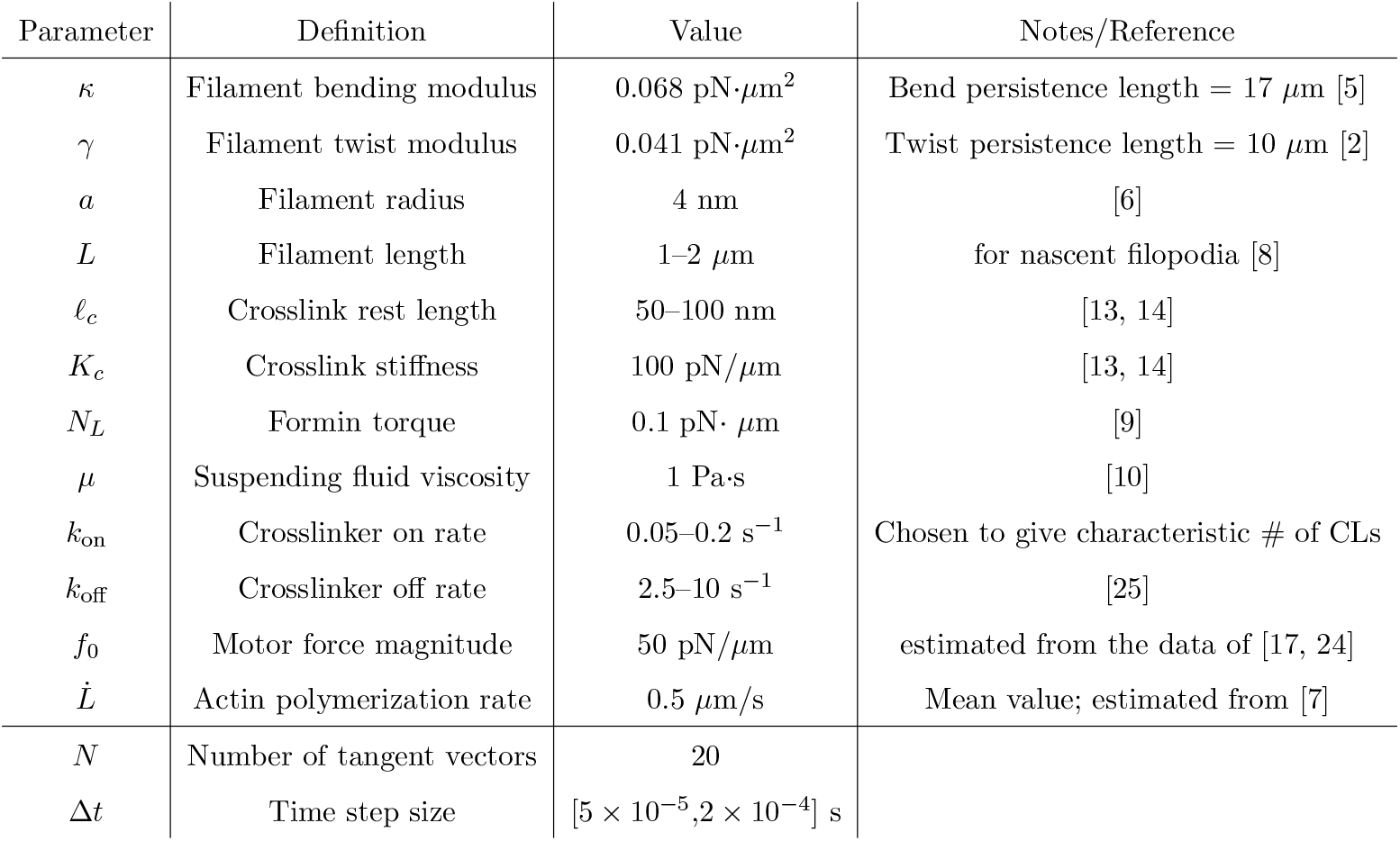
Parameters for our simulation study.

#### Movie legends

Movies of 22-filament bundles with and without motor and twisting activity, showing that motors and twist synergize to generate a compact bundle. Crosslinks are shown in yellow, filaments in white, and formins in magenta. The green dots show the clamped ends, which represent the branched actin network beneath the filopodium. The left side of the movie shows a side view, while the right side shows a top view (in this case, the filaments appear to move outward because of the perspective of the observer).

Movie 1: CCW Motors and CCW formin twisting. Movie 2: No motors, but CCW formin twisting.

Movie 3: CCW Motors, but no formin twisting.

Movie 4: CCW Motors, with CW formin twisting (opposite direction).

---

**Figure S1:**
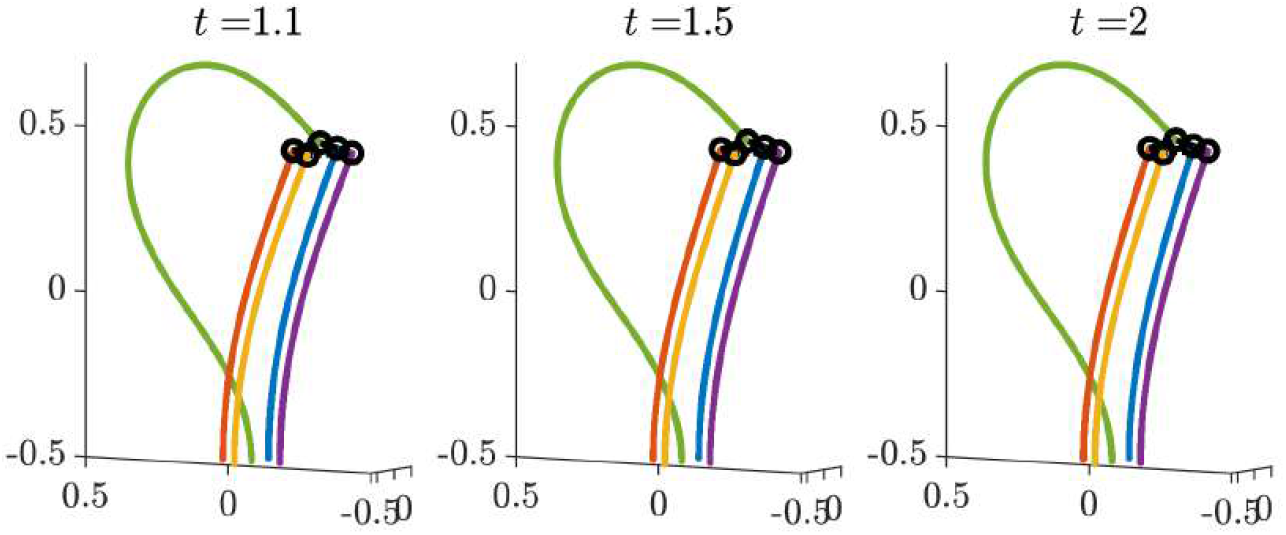
Initial simulation of fiber buckling with polymerization and crosslinking. This is a continuation of Fig. 1C to show relaxation of the central fiber to be slightly above the ends of the others, so that the downward cross linking force balances the upward bending force.

**Figure S2:**
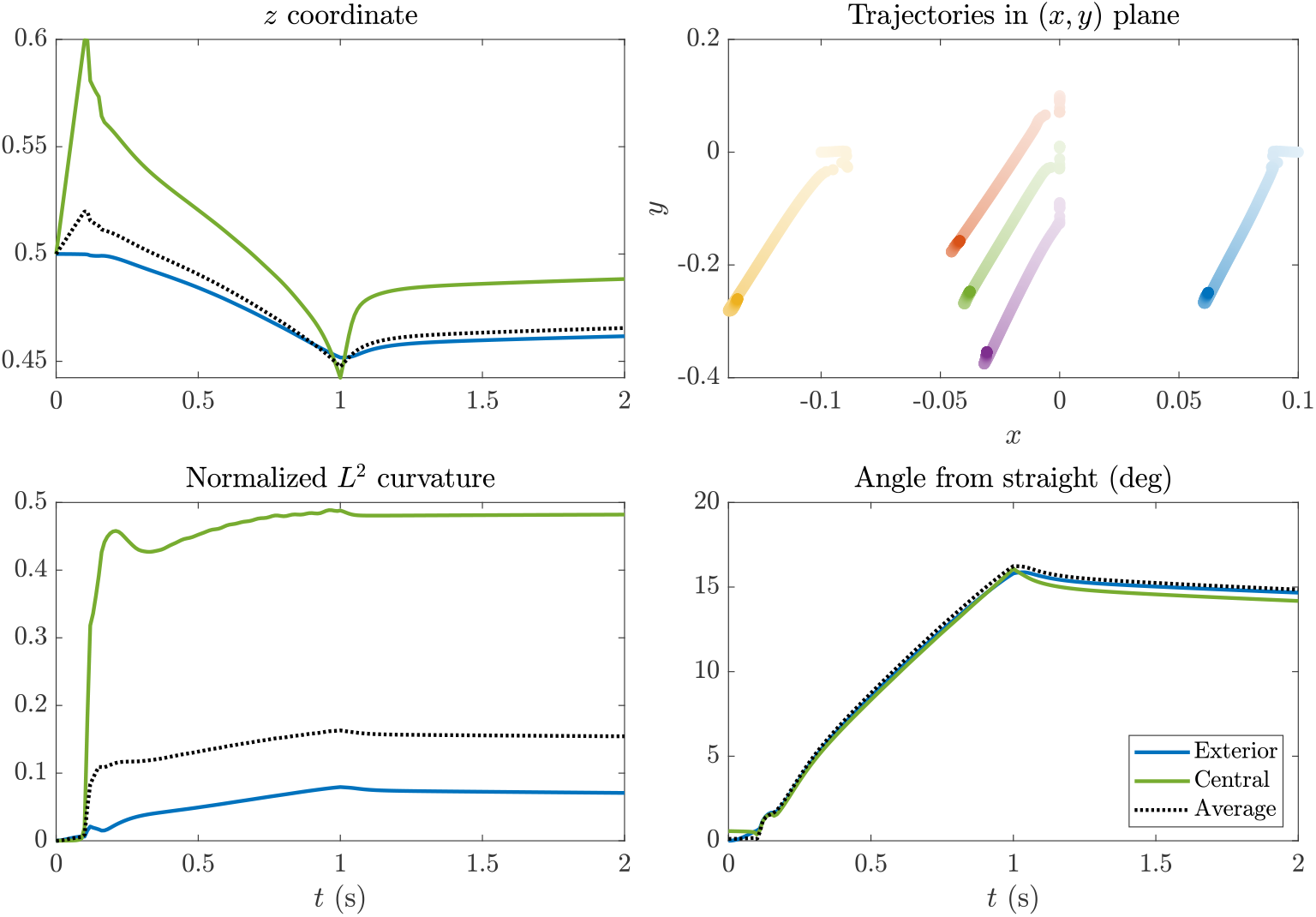
Statistics for the first buckling simulation shown in Figs. 1C and S1. The top right plot shows the trajectories in the (*x, y*) plane, where the colors correspond to the fibers in Figs. 1C and S1, and darker colors denote later times. For the statistics plots, the blue (green) line shows the trajectory of the blue (green) fiber (the blue fiber has the same statistics as the other peripheral fibers by symmetry), while the dotted black line shows the average across the five filaments. Definitions of the *z* coordinate, curvature and “angle from straight” can be found in Section “Quantification of the filopodial bundle’s shape” in the Methods section of the main text.

**Figure S3:**
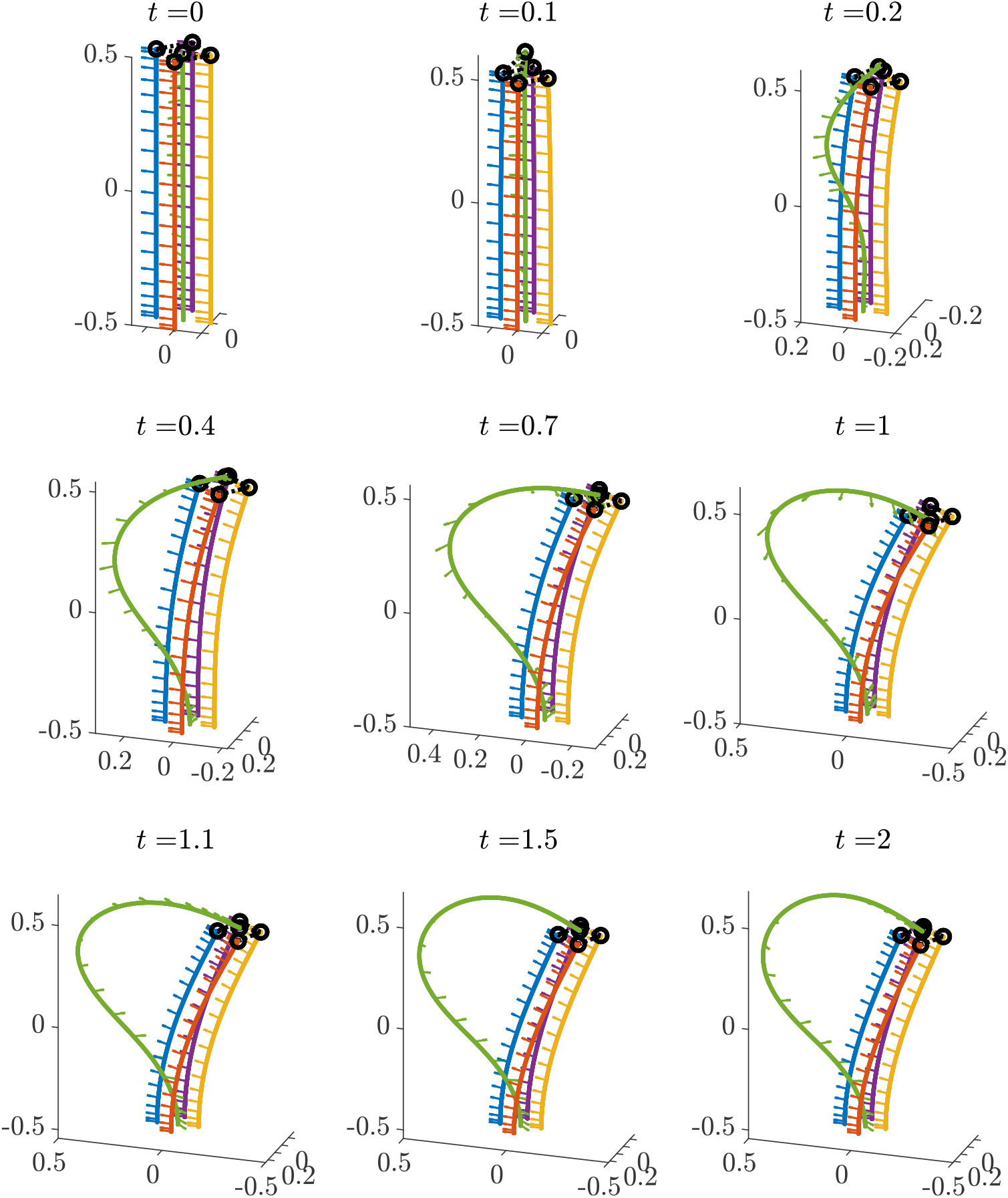
Buckling in a bundle due to polymerization, but with added twist along the fiber centerline. Polymerization occurs only for the middle green filament, with 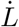 = 1 *μ*m/s on 0 ≤ *t* ≤ 1. Arrows show the material frame vectors.

**Figure S4:**
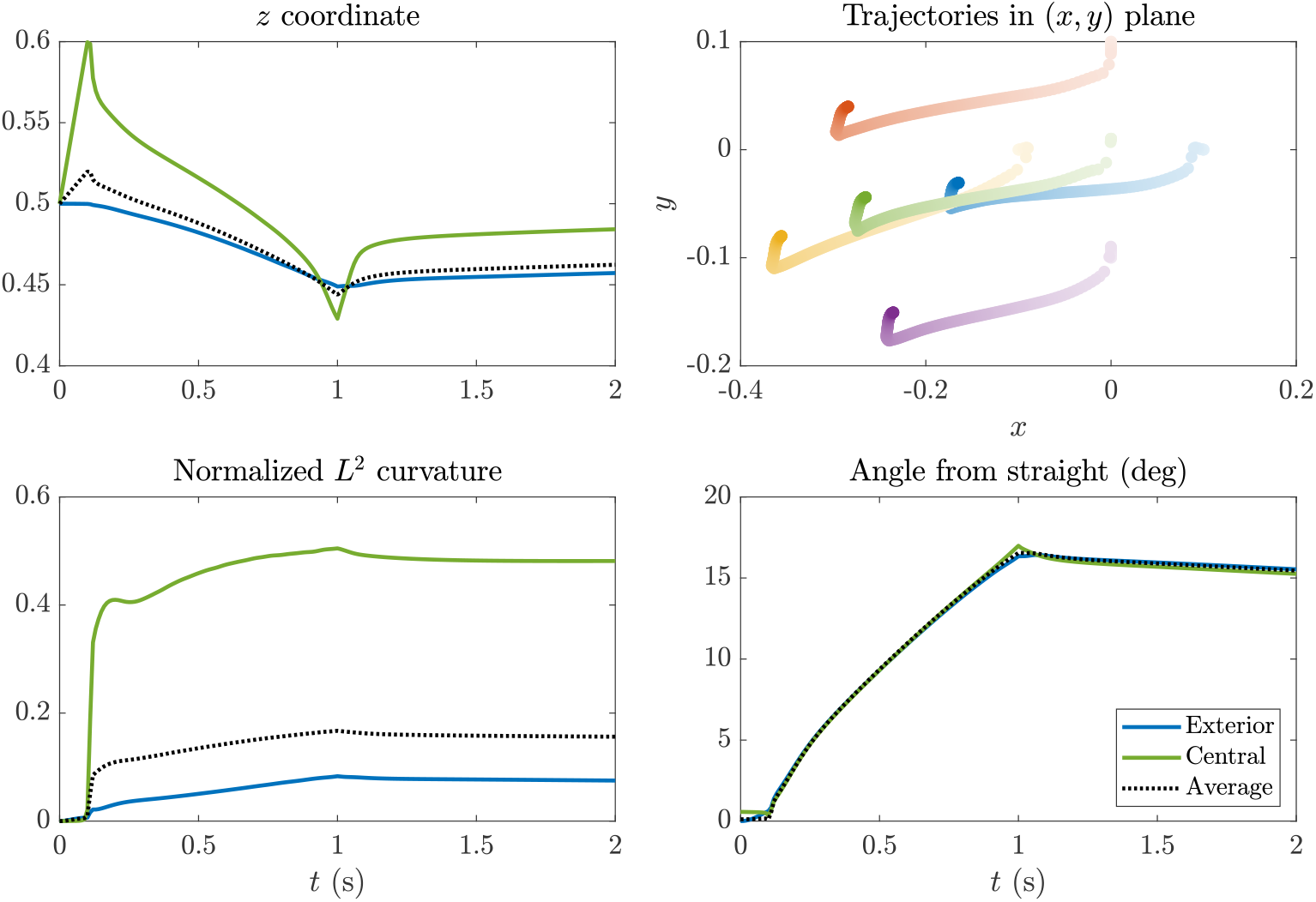
Statistics for the buckling simulation shown in Fig. S3. The top right plot shows the trajectories in the (*x, y*) plane, where the colors correspond to the fibers in Fig. S3, and darker colors denote later times. For the statistics plots, the blue (green) line shows the trajectory of the blue (green) fiber in Fig. S3, while the dotted black line shows the average across the five filaments. Definitions of the *z* coordinate, curvature and “angle from straight” can be found in Section “Quantification of the filopodial bundle’s shape” in the Methods section of the main text.

**Figure S5:**
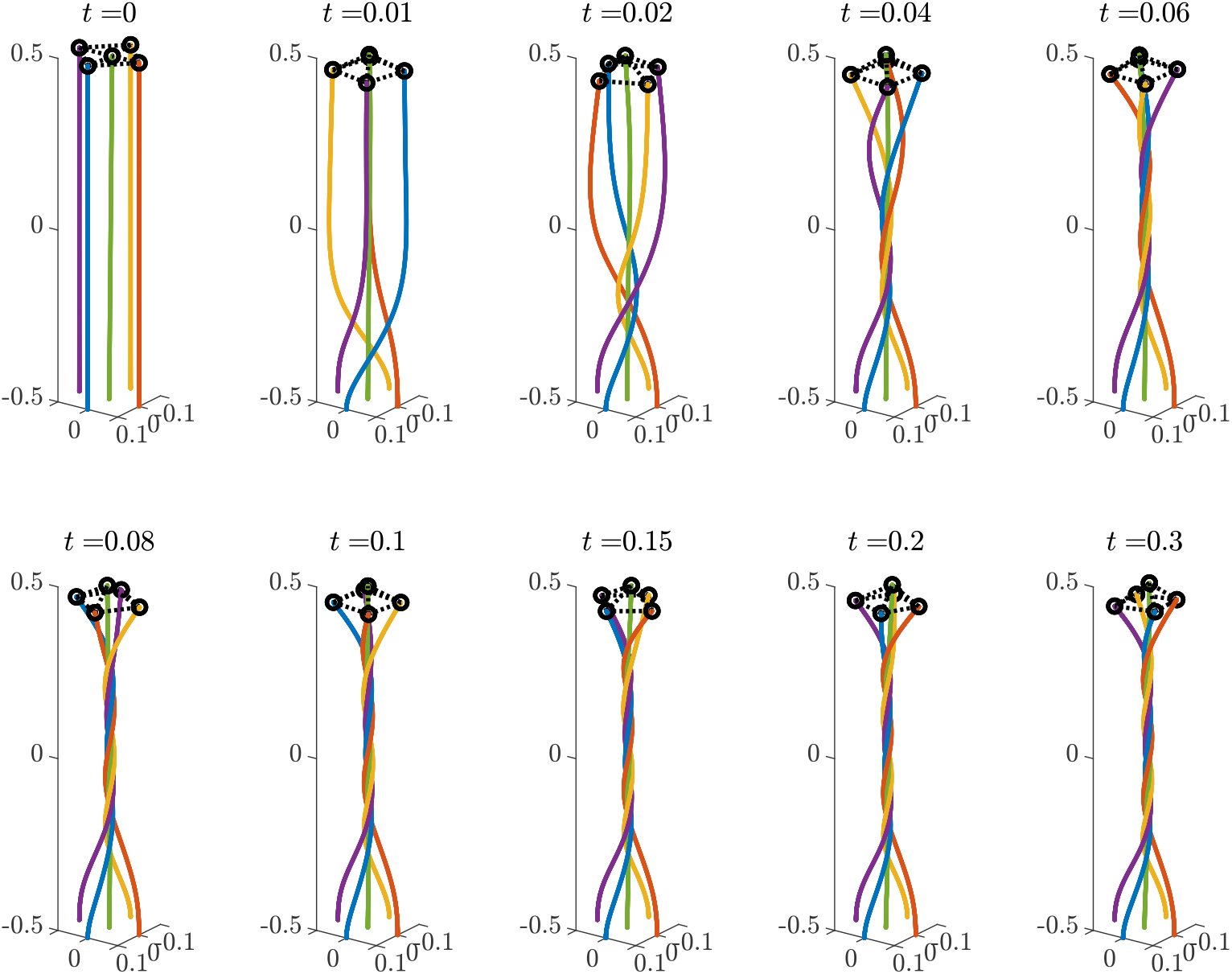
Dynamics of five-filament bundle when the motors act on the edges (outer 1*/*4) of the filopodium and along the whole length of the filaments, (*c*_*m*_ = 1). The tips of the outer filaments are permanently crosslinked (dashed segments) to the tip of the central filament; rings are the endpoints of the crosslinks. Filaments are color-coded for visualization. Time is in seconds, distances are in *μ*m.

**Figure S6:**
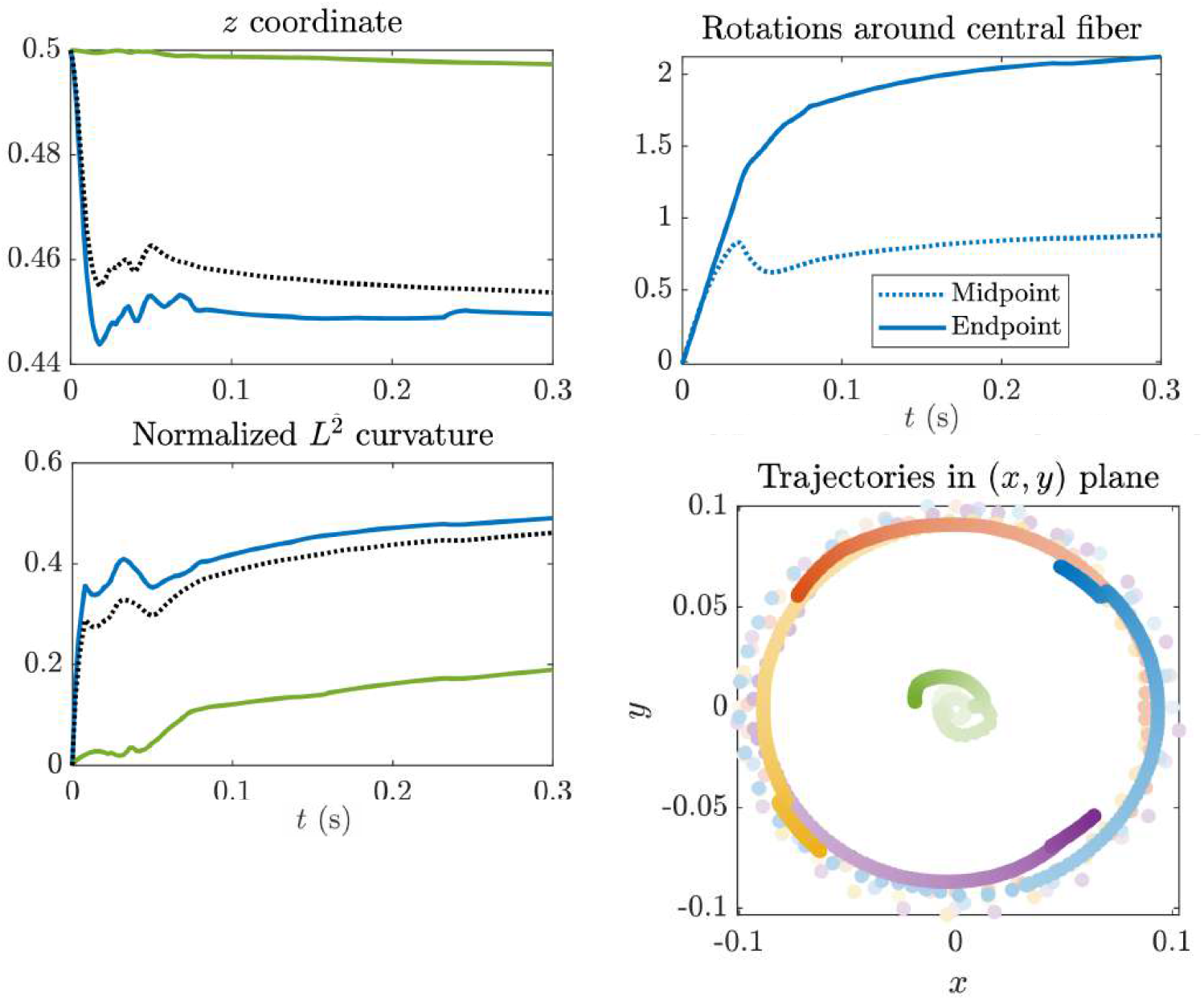
Statistics for the filopodial bundle shown in Fig. S5. Quantities, as functions of time, for the central fiber are in green, for the peripheral fibers – in blue; dashed curves show the averages for all fibers. Top row, left: *z*-coordinates of the filament tips, in *μ*m. Bottom row, left: filaments’ average curvatures (normalized by the curvature of a circle with the same circumference as the filament length). Top row, right: number of rotations of the midpoint (dashed) and endpoint (solid) of the peripheral fiber around the central fiber. Bottom row, right: trajectories of the tips of the filaments projected onto the (*x, y*) plane, color-coded for individual filaments. Light/dark colors correspond to the beginning/end, respectively.

**Figure S7:**
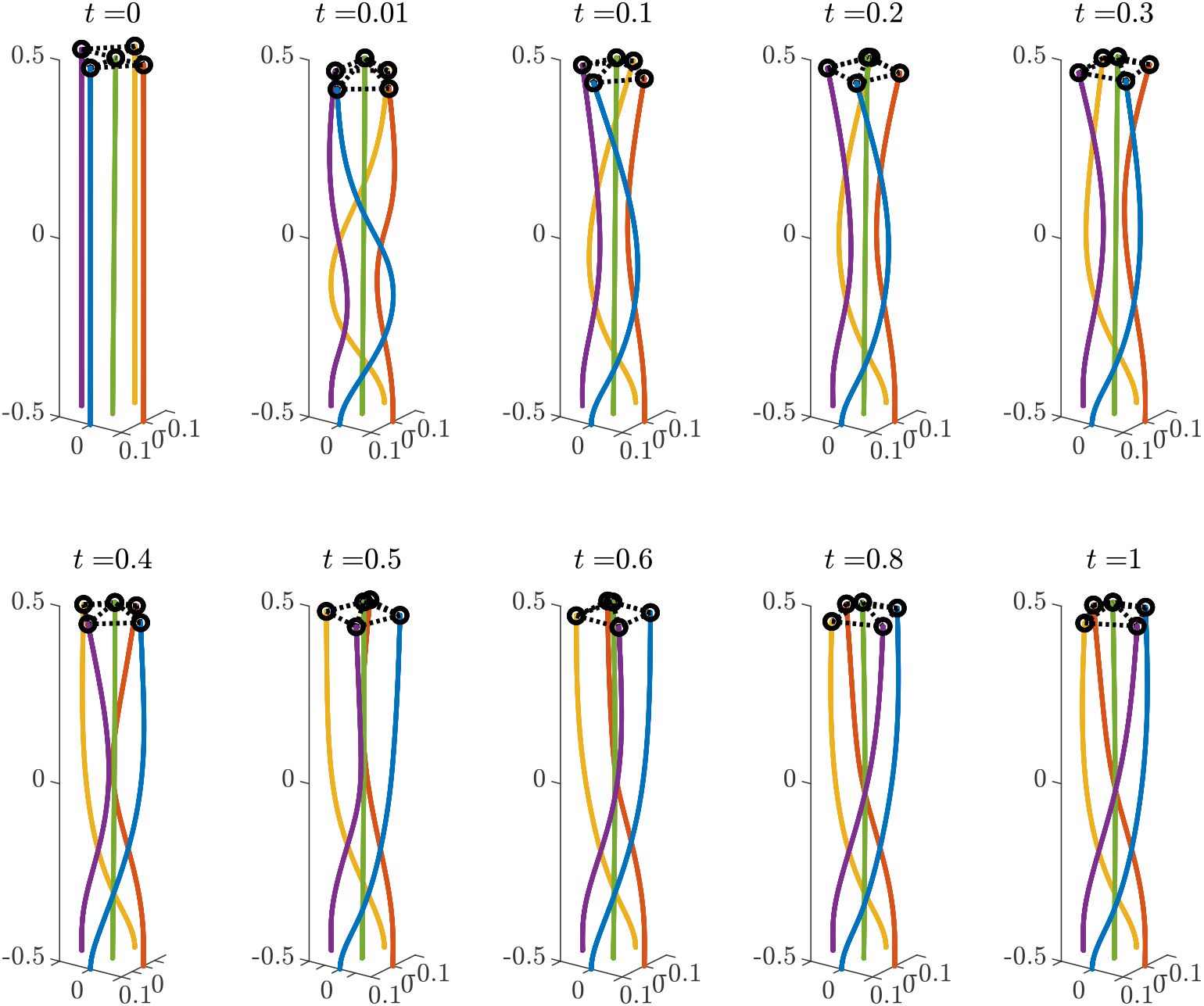
Dynamics of five-filament bundle when the motors act on the edges (outer 1*/*4) of the filopodium and along *half* the length of the fiber (*c*_*m*_ = 1*/*2). The tips of the outer filaments are permanently crosslinked (dashed segments) to the tip of the central filament; rings are the endpoints of the crosslinks. Filaments are color-coded for visualization. Time is in seconds, distances are in *μ*m.

**Figure S8:**
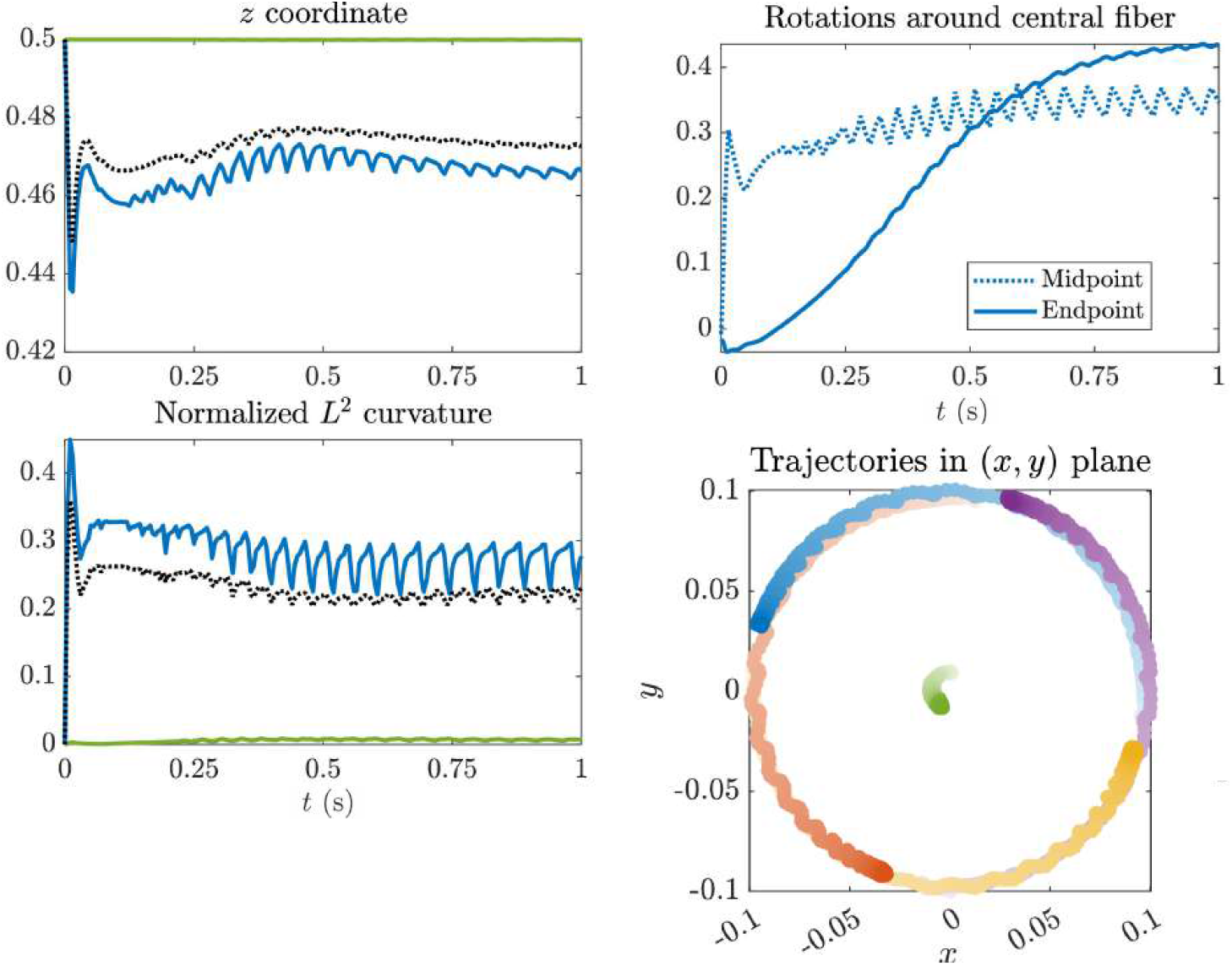
Statistics for the filopodial bundle shown in Fig. S7. Quantities, as functions of time, for the central fiber are in green, for the peripheral fibers – in blue; dashed curves show the averages for all fibers. Top row, left: *z*-coordinates of the filament tips, in *μ*m. Bottom row, left: filaments’ average curvatures (normalized by the curvature of a circle with the same circumference as the filament length). Top row, right: number of rotations of the midpoint (dashed) and endpoint (solid) of the peripheral fiber around the central fiber. Bottom row, right: trajectories of the tips of the filaments projected onto the (*x, y*) plane, color-coded for individual filaments. Light/dark colors correspond to the beginning/end, respectively.

**Figure S9:**
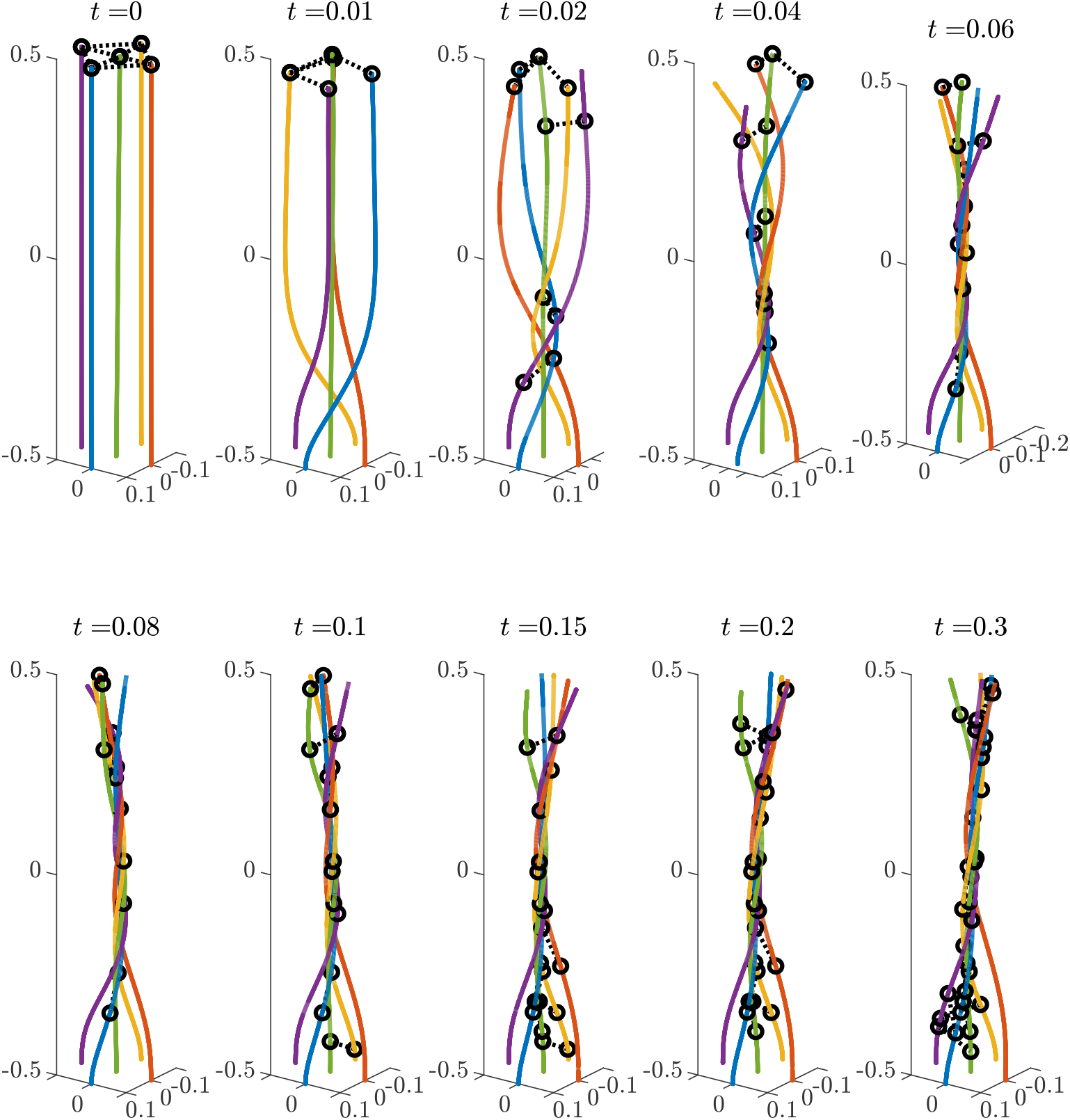
Dynamics of five-filament bundle when the motors act on the edges (outer 1*/*4) of the filopodium and along the whole length of the fiber (*c*_*m*_ = 1). The crosslinks (dashed segments) are transient; rings are the endpoints of the crosslinks. Initially, only the tips of the outer filaments are crosslinked to the tip of the central filament. Filaments are color-coded for visualization. Time is in seconds, distances are in *μ*m.

**Figure S10:**
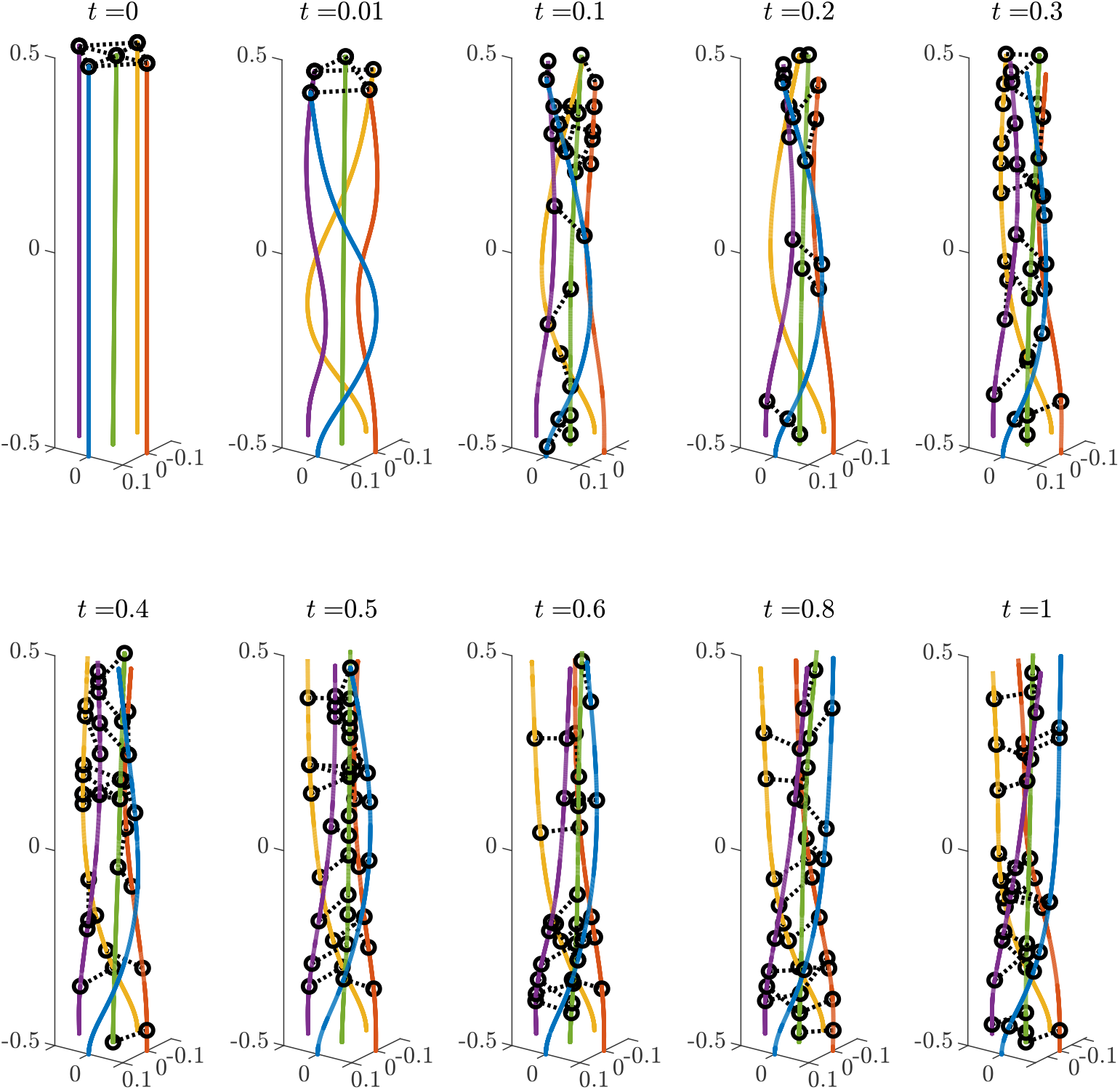
Dynamics of five-filament bundle when the motors act on the edges of the filopodium and along the half length of the fiber (*c*_*m*_ = 1*/*2). The crosslinks (dashed segments) are transient; rings are the endpoints of the crosslinks. Initially, only the tips of the outer filaments are crosslinked to the tip of the central filament. Filaments are color-coded for visualization. Time is in seconds, distances are in *μ*m.

**Figure S11:**
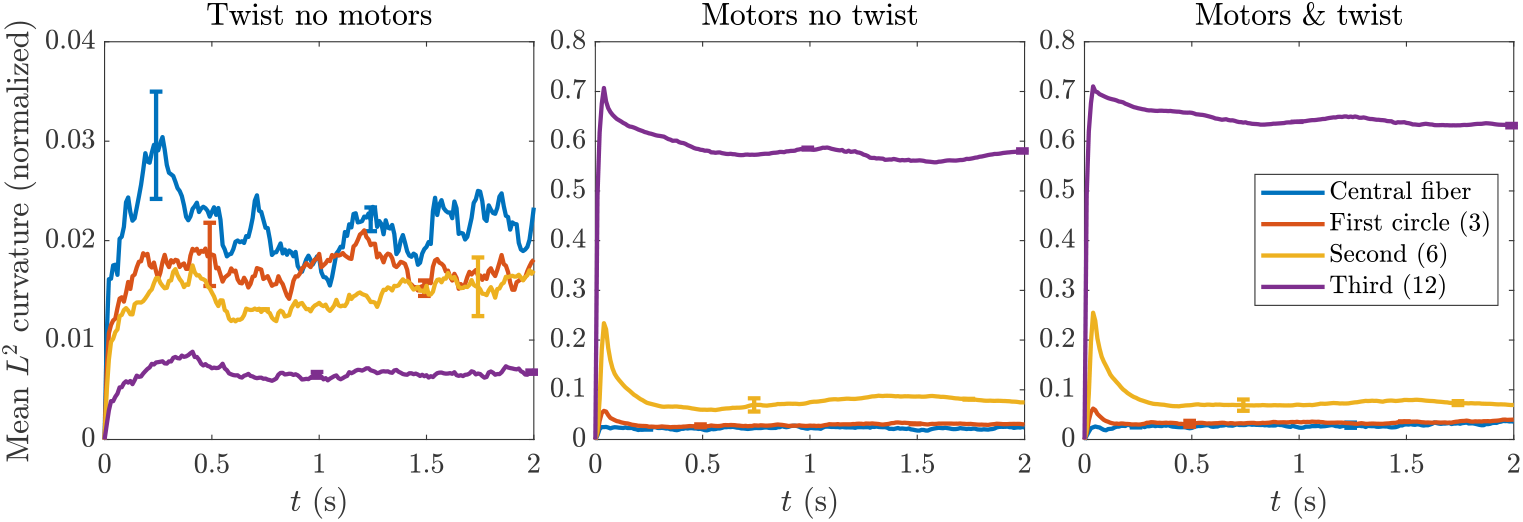
Mean *L*^2^ curvature of filaments, normalized by the curvature of a circle with the same circumference in the 22-filament bundle. From left to right, we show simulations with twist but no motors (Fig. 4B), with motors but no twist (Fig. 4C), and both motors and twist (Fig. 4A), segregating the filaments by what circle their base belongs to.

**Figure S12:**
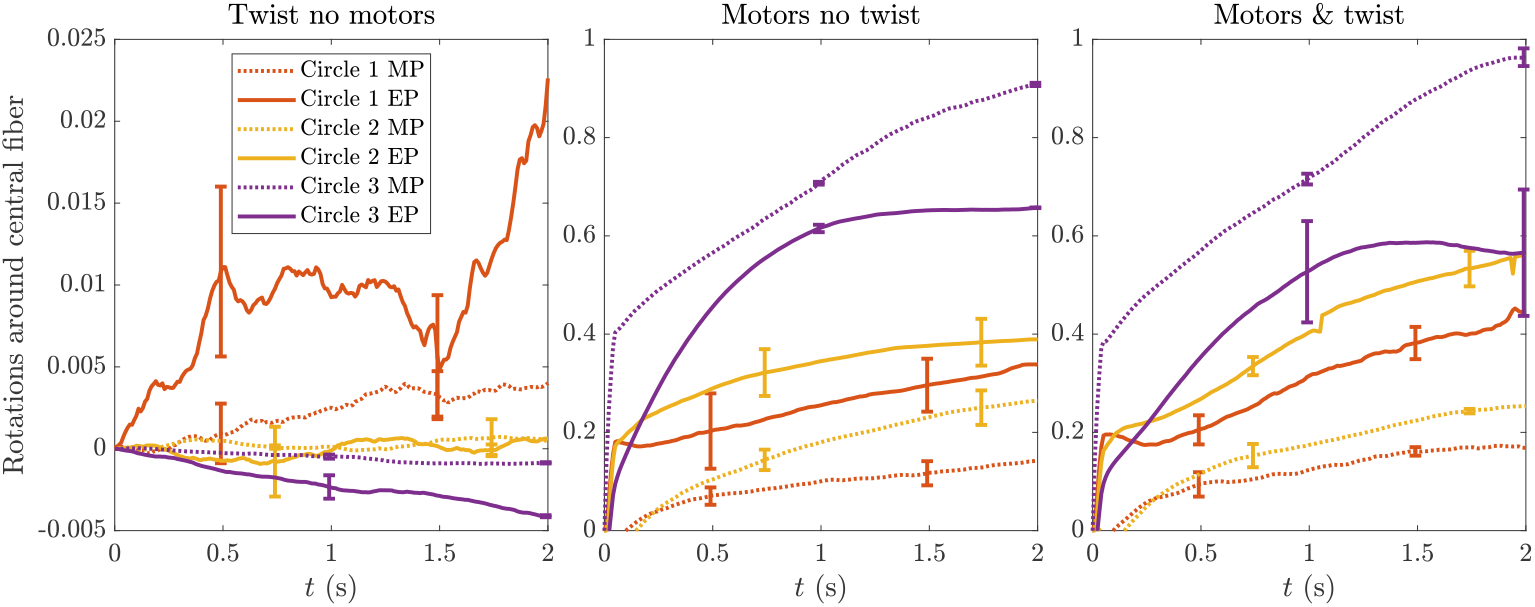
Number of rotations of each circle of filaments around the central filament in the 22-filament bundle. From left to right, we show simulations with twist but no motors (Fig. 4B), with motors but no twist (Fig. 4C), and both motors and twist (Fig. 4A), segregating the filaments by what circle their base belongs to. MP = midpoint, EP = endpoint.

## Notes

### Competing Interest Statement

The authors have declared no competing interest.

